# ROPIUS0: A deep learning-based protocol for protein structure prediction and model selection and its performance in CASP14

**DOI:** 10.1101/2021.06.22.449457

**Authors:** Mindaugas Margelevičius

## Abstract

Protein structure prediction has recently been revolutionized when AlphaFold2 [1] predicted protein structures with near-experimental accuracy in the latest CASP14 season of critical assessment of methods of protein structure prediction (CASP). Among numerous implications, this breakthrough has led to a rapidly growing number of high-quality structural models [2]. We present a protocol ROPIUS0 for protein structure prediction and model selection and discuss its benefits in the new era of structure prediction. At the core of the ROPIUS0 protocol is the deep learning module developed for the selection of protein structural models. It is shown that the direct use of predicted inter-residue distances may be sufficient to discriminate between correct and incorrect protein folds, considering only a small fraction of predicted distances. We extensively tested the protocol: In the latest CASP14 prediction season, a ROPIUS0 variant based on model selection ranked 13th in the category of tertiary structure prediction. Its performance is on par with top-performing automated prediction servers when tested on the CASP13 dataset, and it performs similarly on a CAMEO dataset. The results suggest ways to improve searching for structurally similar and homologous proteins without considerably increasing speed. Our new open-source threading tool based on comparing a subset of inter-residue distances demonstrates the effectiveness and application of the deep learning module of the ROPIUS0 protocol.

## 1 Introduction

The problem of protein three-dimensional (3D) structure prediction is fundamental to Bioinformatics, and structure prediction is particularly important in biomedical research. Knowledge of the structure and thermodynamics of the protein can lead to a much deeper understanding of biological processes. It would greatly facilitate vaccine and drug design [3].

Before the emergence of the AlphaFold2 method [1], the common classification distinguished between two basic approaches to protein structure prediction [4]. One is based on modeling by homology to solved structures [5]. The sequence of the protein of interest, the target, is searched for similarities in a database of sequences with known 3D structure or structures using homology search [6], [7] or threading tools [8], [9]. A significant similarity to a protein from the database suggests that the target protein can adopt a similar structure, and the identified protein can represent a good candidate for using it as a template to model the structure of the target protein. The accuracy of a structural model strongly depends on the extent of the template’s structural similarity to the target protein and the alignment accuracy between the template and the target. Modeling by homology remains an important approach to structure prediction, as it represents a computationally cheap alternative to de novo structure prediction.

De novo structure prediction is the other approach, which predicts the structure without using structural templates. De novo structure prediction has improved dramatically over the past decade [10], with the most successful methods being those that use predicted distances or contacts between residue pairs as restraints [11], [12], [13]. (AlphaFold2 does not use distance predictions explicitly but rather models pair representations.) Restraints are typically generated using artificial neural networks (NNs). A standard choice for the NN backbone architecture has been a residual network (ResNet) [14] due to its good learning and performance characteristics and convolutional nature, which fits well with learning on variable-length proteins. Given the type of NN architecture, the set of input data to the NN, the NN depth expressed in hidden layers, training policy and the size of the training dataset are among the most influential factors that affect the accuracy of predictions.

Protein modeling is accompanied by model accuracy estimation or model selection. Its goal is to select from the set of models generated during the modeling process models structurally most similar to the target protein. More generally, given the target sequence and a set of structural models, a model selection algorithm aims to select the most accurate model, and, therefore, it is as important as discriminating between correct and incorrect protein folds. In the context of independent model accuracy estimation [15], [16], the progress in deep learning has fostered the application of deep NNs to predicting model accuracy locally for each residue [17], [18].

The present study introduces a protocol ROPIUS0 (an acronym for “Restraint-Oriented Protocol for Inference and Understanding of protein Structures”) for protein structure prediction and model selection. The rationale behind the ROPIUS0 protocol is to apply homology modeling when structural templates can be identified. Otherwise, it is aimed at selecting the most accurate models from a set of independently generated structural models.

Model selection is expected to be accurate if it produces estimates that correlate with discrepancies between the model and the target. According to the ROPIUS0 protocol, predicted distances between residues are contrasted with the distances observed in the model, and the differences are used to calculate model quality scores. This way, the accuracy of predictions determines the accuracy of model selection. Given the great prediction power of deep learning models, the deep learning module is at the core of the ROPIUS0 protocol.

ROPIUS0 employs a convolutional encoder-decoder NN architecture for predicting the distribution of distances between residues. The encoder part encodes the input into a latent space capturing dependencies in every input receptive field patch. The decoder part produces distance predictions by operating on the information encoded by the encoder. Predictions are therefore a result of considering the input representing the entire target.

The present study introduces a protocol ROPIUS0 (an acronym for “Restraint-Oriented Protocol for Inference and Understanding of protein Structures”) for protein structure prediction and model selection. The rationale behind the ROPIUS0 protocol is to apply homology modeling when structural templates can be identified. Otherwise, it is aimed at selecting the most accurate models from a set of independently generated structural models.

Model selection is expected to be accurate if it produces estimates that correlate with discrepancies between the model and the target. According to the ROPIUS0 protocol, predicted distances between residues are contrasted with the distances observed in the model, and the differences are used to calculate model quality scores. This way, the accuracy of predictions determines the accuracy of model selection. Given the great prediction power of deep learning models, the deep learning module is at the core of the ROPIUS0 protocol.

ROPIUS0 employs a convolutional encoder-decoder NN architecture for predicting the distribution of distances between residues. The encoder part encodes the input into a latent space capturing dependencies in every input receptive field patch. The decoder part produces distance predictions by operating on the information encoded by the encoder. Predictions are therefore a result of considering the input representing the entire target.

The main difference from existing methods that use predicted distances between residues to predict model quality scores [19] is that ROPIUS0 uses distances predicted by the deep learning module directly for model accuracy estimation and selection. We show that the direct use of predicted distances may be sufficient to achieve satisfactory results even in the presence of some limitations, which we discuss in Section 3. We registered and tested ROPIUS0 (ropius0) and its variant ROPIUS0QA (ropius0QA) in the latest 14th CASP [10] season. ROPIUS0QA was officially ranked 13 (and among the top 10 independent groups) in the category of tertiary structure prediction. When tested on the CASP13 dataset, ROPIUS0QA was found to perform on par with top-performing automated prediction servers. And it performs similarly on a CAMEO dataset.

In this study, we focus on the model selection algorithm since the model selection-based variant ROPIUS0QA produced better results but describe the complete ROPIUS0 protocol for consistency. We further discuss the implications of this algorithm for improving homology search and modeling.

## 2 Results

In this section, we first describe the workflow of the ROPIUS0 protocol and define how ROPIUS0QA differs from it. We then proceed with the results of the protocols.

### 2.1 ROPIUS0 workflow

The goal of the ROPIUS0 protocol is to predict the structure of a given target protein sequence (Fig. 1). The target is modeled by homology modeling, or its structure is selected from independently generated models using the deep learning module (the feature prediction part in Fig. 1). The independently generated models represent CASP-hosted server predictions, which are available for (human) groups assigned three weeks to build models for a target. In general, they represent structural models generated by one or more methods, e.g., web servers.

**Fig. 1.**
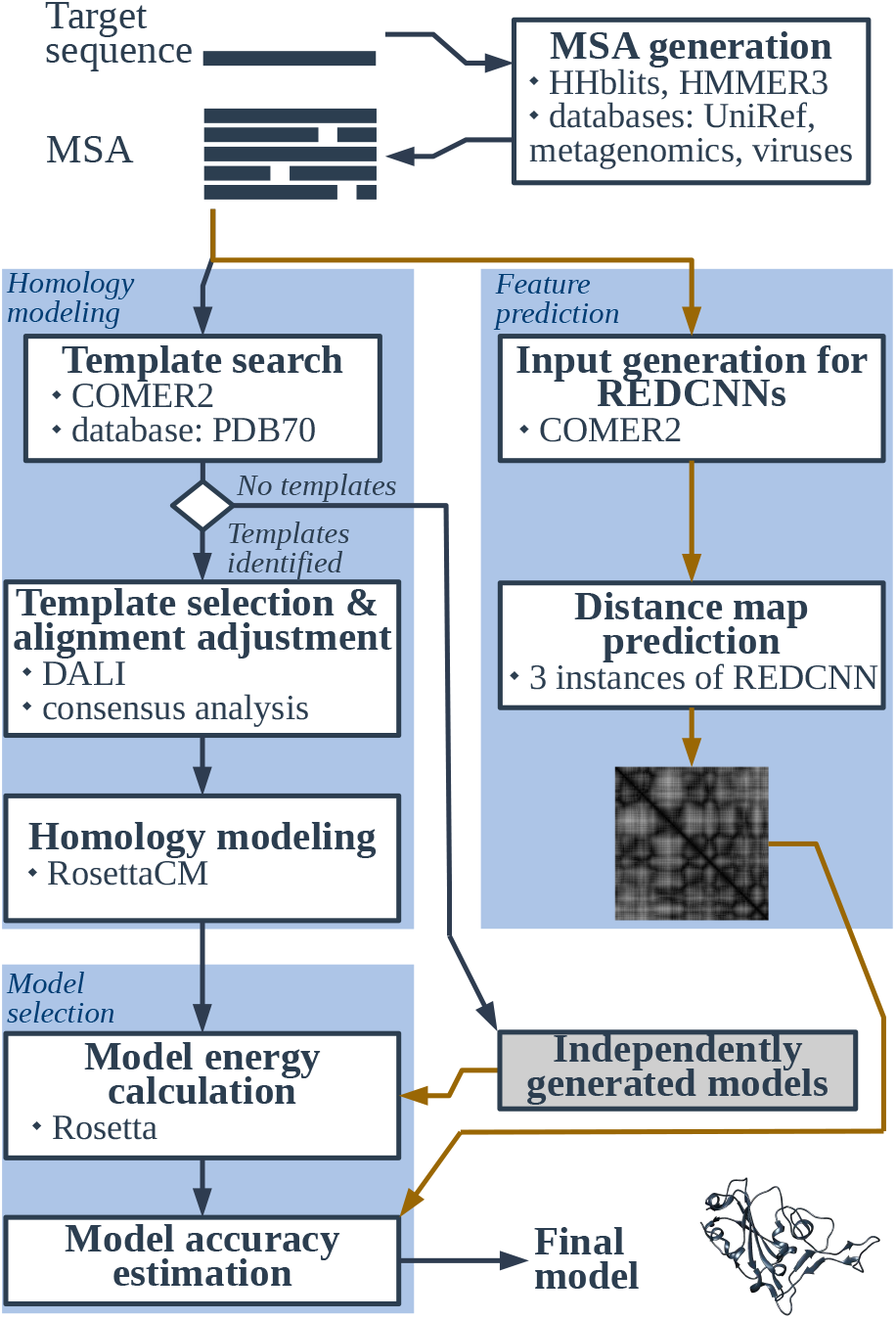
Workflow of the ROPIUS0 protocol. White boxes represent processes. The external and specifically developed tools and resources used in a process are listed at the bottom of the box. The path highlighted in yellow denotes the part of the workflow executed by ROPIUS0QA.

The approach of modeling is determined by the availability of structural templates for homology modeling. Available templates lead to invoking a structure-based procedure for template selection and alignment adjustment, followed by homology modeling. Concurrently, structural features of the target are predicted that are used as restraints for model ranking and selection. The restraints on the distances between the CB atoms of the target protein are generated using residual encoder-decoder convolutional neural networks (REDCNNs).

Structural models generated by either of the modeling approaches are ranked by the model accuracy estimation module, which calculates the energy of each model structure and evaluates the closeness of the distances between CB atoms (CA for Gly) in a model to those predicted by the REDCNNs. The model with the highest estimated accuracy is considered the most probable. The ROPIUS0 components are described in more detail in Section 5.

ROPIUS0QA is a special case of the ROPIUS0 protocol. It is designed to always perform model selection on a set of independently generated models. The difference between ROPIUS0 and ROPIUS0QA lies, therefore, in homology modeling and refinement (Section 5.4). ROPIUS0 generates models by homology and ranks them using the REDCNNs when templates can be identified, whereas ROPIUS0QA always selects among independently generated models. The ROPIUS0QA path in the ROPIUS0 workflow is highlighted in yellow in Figure 1.

### 2.2 Performance in CASP14

CASP is an initiative to rigorously evaluate predictions submitted by registered prediction groups [10]. CASP prediction seasons are organized biennially. The latest CASP14 season for tertiary structure prediction started on 18th May and lasted until 21st August 2020. The sequences of unpublished target structures were issued every workday with a time frame of three days and three weeks for the server and human groups, respectively, to submit their predictions. The human groups were allowed to use the models submitted by the server groups.

At the end of CASP14, the independent assessors evaluated the predictions on a protein domain basis so that the orientation of the domains in a model did not influence the evaluation results. The evaluation takes into account the complexity of each target and is based on calculating the *z*-score for the GDT_TS score [20] of the superposition between a model and the target domain. The total *z*-score accumulated over all domains determines a group’s position in the final ranking [4].

We registered ROPIUS0 and ROPIUS0QA for the CASP14 season as two human prediction groups (group IDs 254 and 039, respectively). The results are shown in Figure 2. The official ranking for tertiary structure prediction is available in Ref. [21].

**Fig. 2.**
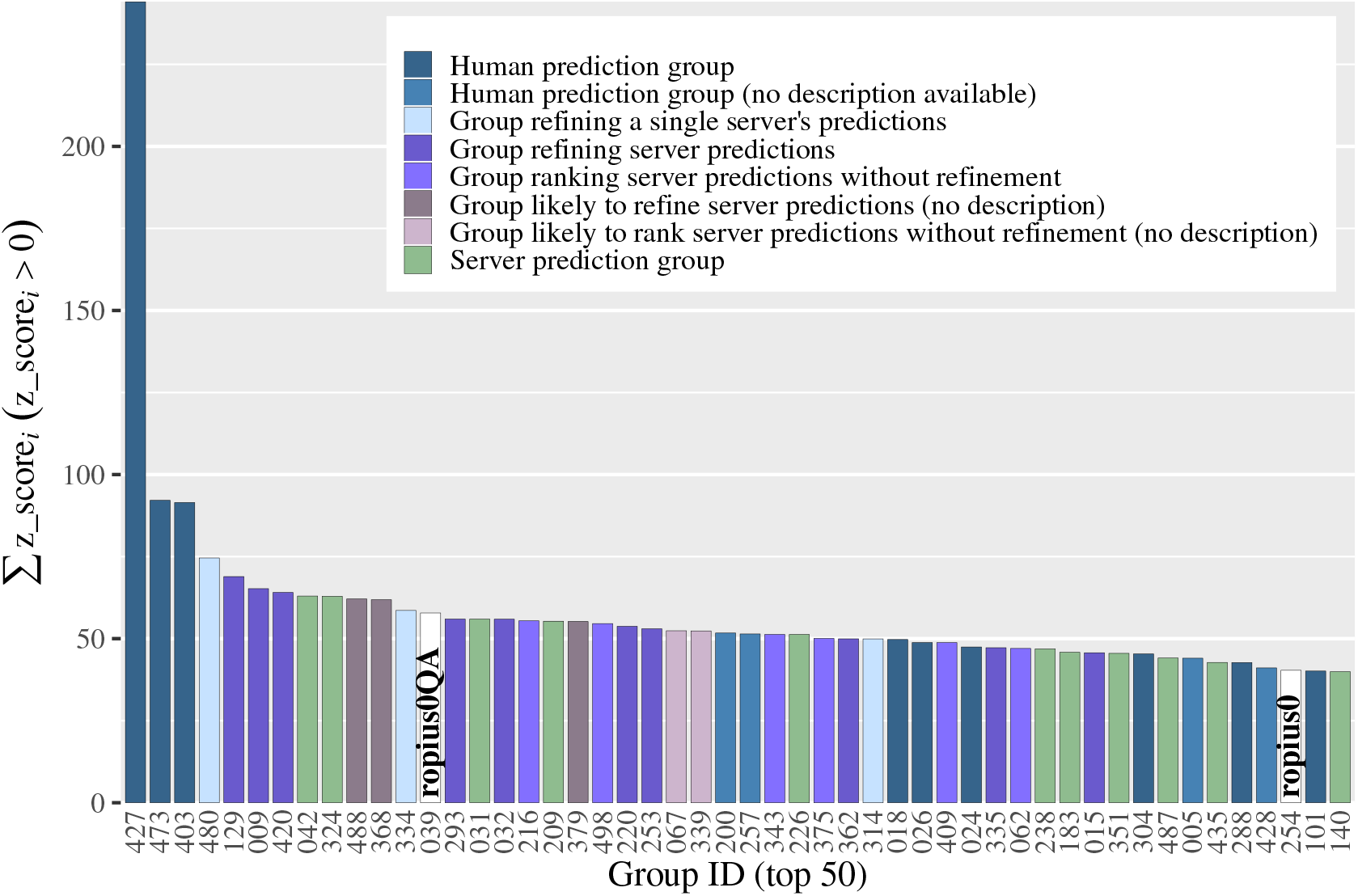
Ranking of the groups registered for the CASP14 season in the tertiary structure prediction category. The ranking is based on the sum of the *z*-scores found to be positive for the structural models designated by the group under consideration as best (1st model) [21]. The top 50 groups are shown. Server prediction groups are represented in green.

The first observation is that one group, AlphaFold2 (427), stands out above the rest, with most of the targets modeled with high accuracy. Evidently, the group’s deep learning approach [1] has revolutionized protein structure prediction.

ROPIUS0 and ROPIUS0QA rank 48th and 13th, respectively, among 146 groups. The ROPIUS0QA approach of using model selection from the server predictions without any further refinement was more accurate than the ROPIUS0 approach including homology modeling.

We see three reasons for this effect. The first is that protein structure prediction methods have improved substantially over recent years. Deep learning frameworks are able to accurately predict structural characteristics for using them as restraints in protein structure optimization and lead to high-quality models without considering templates. Second, the top-performing groups do not use homology modeling directly (except for group 183, tFold-CaT, among the top 50 groups). A typical approach instead converts structural information of identified templates to additional restraints for optimization or uses them as additional input for a more confident prediction of target inter-residue distances and orientations. Its advantage in modeling insertions in the target sequence is evident. Yet, it can also be seen from the perspective of processing deletions. An attempt to remove a fragment in the template structure by homology modeling can end up (to the extent depending on the modeling tool) with the distorted geometry of aligned fragments, even though the alignment can be correct. The third reason is model selection. The model accuracy estimation module was mainly tested through the ROPIUS0QA path. This module for ranking homology models was used by the ROPIUS0 protocol only in the second half of the CASP14 season (Section 5.4). The results show that using it during the entire CASP14 season would have been advantageous.

Judging from the available description of the groups’ methods [22], ROPIUS0QA ranks best among the groups that made selections from the server predictions and submitted models without refinement (Fig. 2). Although ROPIUS0QA ranks models by providing accuracy estimates representing errors in Angstroms, no efforts were made for these estimates to correlate well with GDT_TS scores. The primary goal, instead, was to select the best available model (Section 5.8).

ROPIUS0 was also registered in the model accuracy estimation (EMA) category. Model ranking in this category differed from the ranking by ROPIUS0QA to make selected models more consistent with the ROPIUS0 predictions (Section 5.4).

The rest of the Results section, therefore, focuses on the analysis of the ROPIUS0QA method and model selection, while Section 5 and Supplementary Sections S1.1–S1.3 (Supplementary Figs. S1–S4) present and provide results of a structure-based sequence alignment algorithm for improving alignment accuracy.

### 2.3 Test results on the CASP13 targets

To further explore the ROPIUS0QA’s ability to correctly select structural models, we tested ROPIUS0QA on the CASP13 dataset. The test was performed in the same manner as that used during the CASP14 season. ROPIUS0QA was applied to the full targets, and the models selected from the CASP-hosted server predictions were evaluated on a target domain basis, following the (same) *z*-score-based official procedure for ranking groups [23].

Normally, ROPIUS0QA uses two sets of thresholds for two application modes that depend on target difficulty (Section 5.8). Figure 3 shows that both ROPIUS0QA and its single-mode version (ropius0QA_SM), which simplify application, achieve similar results, which are on par with those of the top-performing server groups. The conclusion does not change (Supplementary Fig. S5) if the analysis excludes ten targets used to determine the REDCNN application thresholds and five targets/domains having a HMMER [24] hit at a significance threshold of E-value = 0.01 to a member of the REDCNN training and validation datasets. (A significance threshold of E-value = 0.01 for searching the sequences of the training and validation datasets corresponds to E-value ≈ 330 for searching a UniRef50 sequence database.)

**Fig. 3.**
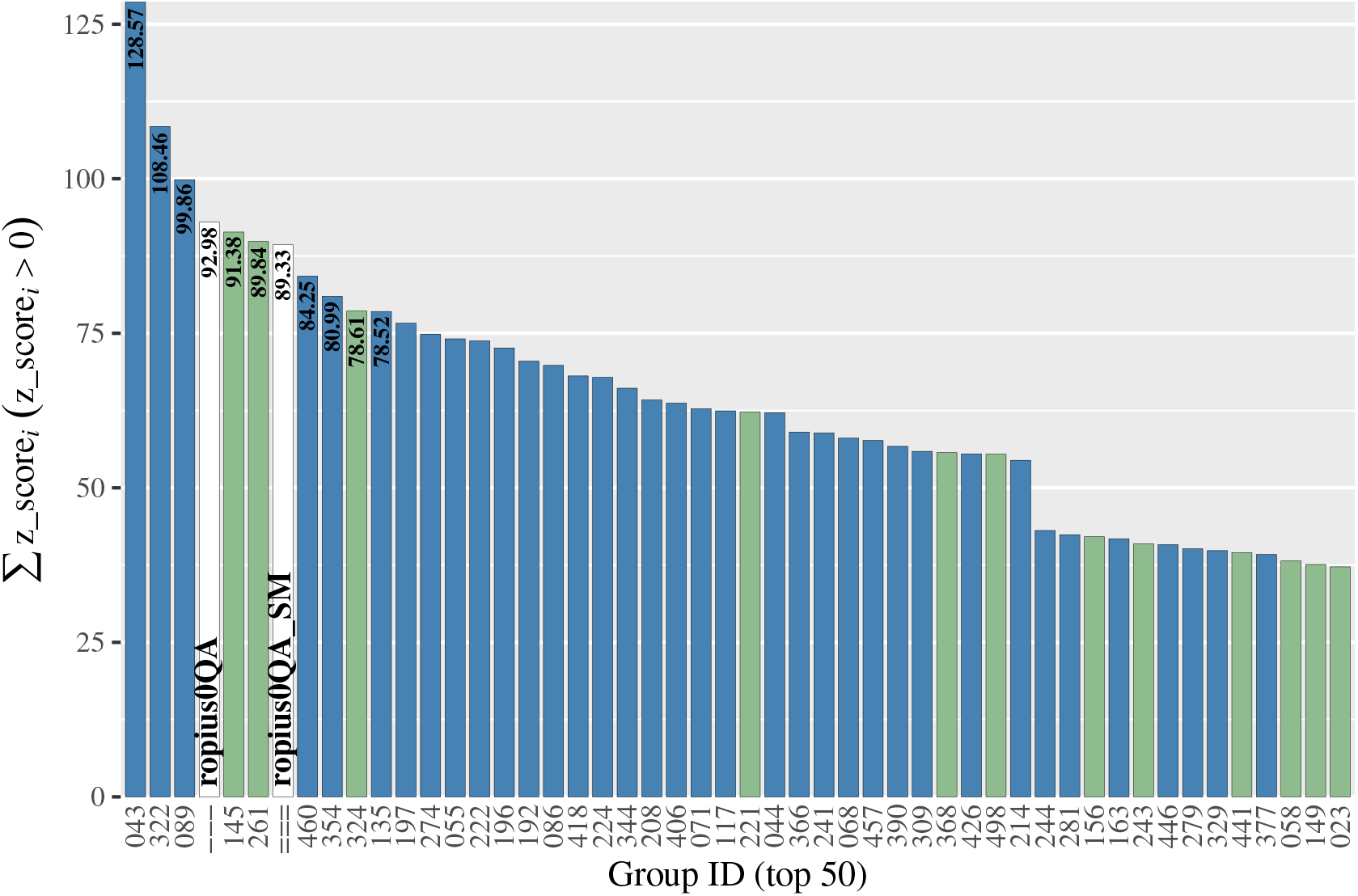
Results of the test on the CASP13 targets (104 domains) in the tertiary structure prediction category. ROPIUS0QA and ROPIUS0QA_SM were not registered for the CASP13 season. The ranking is based on the sum of the *z*-scores found to be positive for the structural models designated by the group under consideration as best (1st model) [23]. The top 50 groups are shown. Server prediction groups are represented in green.

Table 1 and 2 report ROPIUS0QA’s performance contrasted with DeepAccNet [25] and two of the best single-model accuracy prediction methods in CASP13 [26]: ProQ3D [27] and QMEANDisCo (FaeNNz) [16] evaluated using latest sequence databases in the same way as ROPIUS0QA (see Supplementary Section S1.4 for details). Importantly, ROPIUS0QA compares favorably even when multiple sequence alignments (MSAs) for generating input to the REDCNNs originated from searching a database (Uni-clust30 [28] 10/2017) with a tool (HHblits [29] v3.0.0), both released before the start of the CASP13 season. The result of this test, which brings ROPIUS0QA closer to the conditions at the time of CASP13, implies that sequence quantity alone does not determine MSA quality. The accuracy and relevance of sequence alignments are just as important.

**TABLE 1.** Evaluation results on the CASP13 targets. An asterisk (*) denotes a ROPIUS0QA version provided only with the Uniclust30 database released before the CASP13 season. The average runtimes per target (Runtime) exclude the time spent on sequence search. The runtimes for each CASP13 target are plotted in Supplementary Figure S6. The CASP13 dataset of the available DeepAccNet predictions [25] lacks data for three targets T0954, TO999, and T1005. The sum of the *z*-scores for three DeepAccNet versions (Standard, Bert, and MSA) excluding these three targets (six domains) is 57.52, 61.19, and 77.97, respectively, s.d., standard deviation.

| Method | $\sum_{z\_score_i > 0} z\_score_i$ | Runtime (s), (s.d.) |
| --- | --- | --- |
| ROPIUS0QA | 92.98 | 133.35 (128.75) |
| ROPIUS0QA_SM | 89.33 | —" |
| ROPIUS0QA* | 90.87 | —" |
| ROPIUS0QA_SM* | 88.47 | —" |
| QMEANDisCo | 65.36 | 58.27 (47.62) |
| ProQ3D | 60.67 | 6104.60 (6635.30) |

**TABLE 2.** Evaluation results on 76 CASP13 target domains. Ten targets used to determine thresholds for the REDCNNs are excluded. Five more targets/domains (T0954, T0966, T0983, T0999, and T1017s1-D1) having sequence similarity to a member of the training and validation sets are excluded too. An asterisk (*) denotes a ROPIUS0QA version using the Uniclust30 database released before the CASP13 season. The results for three DeepAccNet versions (Standard, Bert, and MSA) were obtained from the available predictions on the CASP13 dataset [25].

| Method | $\sum_{z\_score_i > 0} z\_score_i$ |
| --- | --- |
| ROPIUS0QA | 69.77 |
| ROPIUS0QA_SM | 68.26 |
| ROPIUS0QA* | 67.14 |
| ROPIUS0QA_SM* | 66.31 |
| DeepAccNet-MSA | 63.25 |
| DeepAccNet-Standard | 48.69 |
| QMEANDisCo | 48.49 |
| ProQ3D | 47.69 |
| DeepAccNet-Bert | 47.11 |

This is illustrated with two examples (Table 3). ROPIUS0QA’s performance was measured for different variants of input MSAs constructed for the same target. Appending the initial MSA with aligned sequences simulated using the same MSA model [30] does not change the result. In contrast, sequences simulated with noise deteriorate the quality of the MSA and, consequently, the result. The same effect is observed when the number of HHblits iterations for T1005 increases from 2 to 6. The MSA obtained after 6 iterations inflated with weakly related sequences, where the sequences at the significance limit at iteration 2 became highly significant, and the statistical significance of the highly significant sequences at iteration 2 considerably decreased.

**TABLE 3.** Example of the importance of alignment quality rather than quantity alone. The effective number of sequences (Neff) in the MSA and the resultant z_score for different variants of MSAs provided as input to ROPIUS0QA are reported for two CASP13 targets, T1005 and T1015s1. MSE_n*i* represents the MSA obtained after *i* iterations of an HHblits search against the Uniclust30 database. +rndmsa_r0 and +rndmsa_r0.1 represent the original MSA (MSA_n2 or MSA_n4) appended with aligned sequences simulated [30] using the original MSA as a model without noise (r0) and with noise (r0.1), respectively.

|  | T1005 |  |  |  | T1015s1 |  |  |
| --- | --- | --- | --- | --- | --- | --- | --- |
|  | MSA_n2 | +rndmsa_r0 | +rndmsa_r0.1 | MSA_n6 | MSA_n4 | +rndmsa_r0 | +rndmsa_r0.1 |
| Neff | 201.36 | 230.36 | 230.36 | 1992.63 | 51.48 | 100.48 | 100.48 |
| $z\_score$ | 0.73 | 0.73 | 0.06 | -0.07 | 1.73 | 1.73 | -0.72 |

The dependence of ROPIUS0QA’s performance on the amount and diversity of sequences in an MSA for the CASP13 targets is shown in Figure 4. A weak correlation between the effective number of sequences in the MSA (Supplementary Section S2.2) and the distance from the highest GDT_TS score (Pearson’s *r* = −0.13 for Neff, and *r* = −0.22 for log(Neff)) demonstrates ROPIUS0QA’s ability to differentiate between accurate and inaccurate models for a considerable part of the shallow MSAs.

**Fig. 4.**
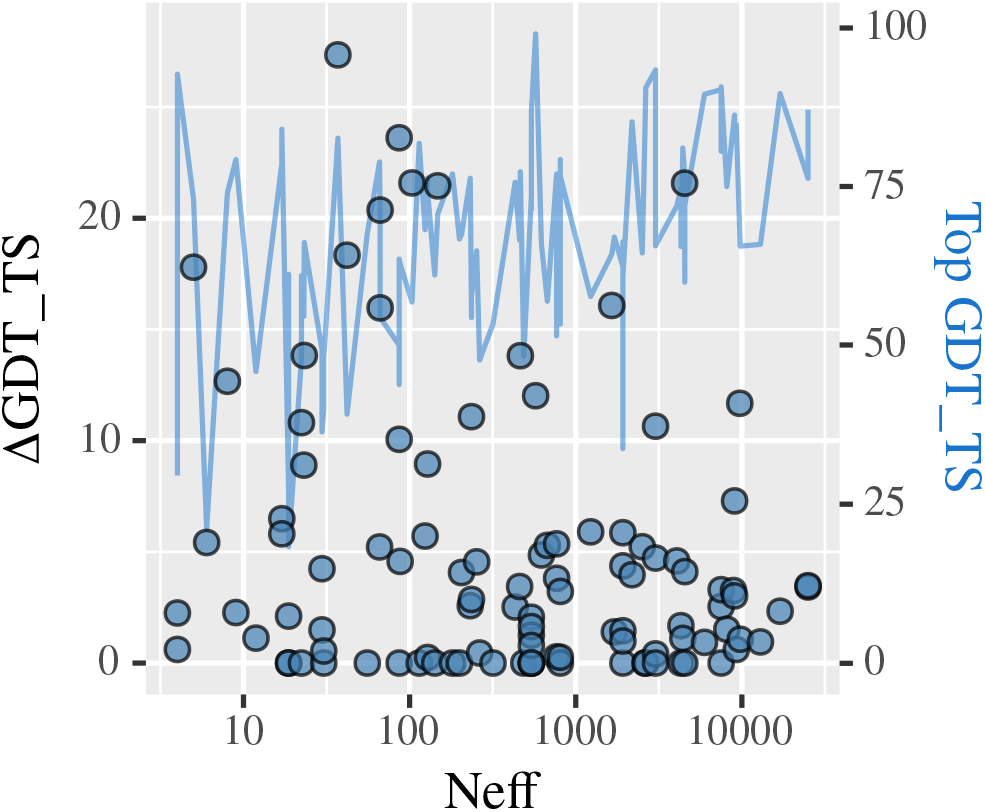
Difference (△GDT_TS) between the highest GDT_TS among the server groups (Top GDT_TS, line) and the ROPIUS0QA GDT_TS plotted against the effective number of sequences (Neff) for the groups’ first models of the CASP13 targets.

### 2.4 Benchmarking on a CAMEO dataset

We also benchmarked ROPIUS0QA on a large dataset of 186 CAMEO [31] targets released over a period of three months from 07/24/2021 through 10/16/2021. ROPIUS0QA was applied to structural models submitted by the registered prediction servers and tested for its ability to select the most accurate model. Under this setting, ROPIUS0QA places 2nd, behind RoseTTAFold [32]. The results are given in Supplementary Section S1.6.

### 2.5 Deep learning performance justification

The REDCNNs in the ROPIUS0 protocol (Fig. 1) were trained on a small set of training data (Supplementary Section S2.1). Yet, their application to model selection has been shown to produce competitive results. An explanation for this phenomenon lies in the application of REDCNN predictions.

Inaccurate distance predictions diminish the ability to select the most accurate structural models. In application, the probability of using inaccurate predictions made by the REDCNNs is reduced, involving only confident predictions. Distance predictions made with probability at least equal to some threshold are retained, while those of lower probability are ignored (Section 5.8).

Figure 5 shows the dependence of the accuracy of distances, predicted by combining the output of the three REDCNNs, on the threshold probability for the CASP14 targets for which the structure is available. (A similar result obtained for the CASP13 targets is shown in Supplementary Fig. S7.) We observe that increasing the threshold probability leads to more accurate inter-residue distance predictions, independently of how far apart residues are in the sequence. Yet, the number of distances predicted with at least the threshold probability decreases with increasing threshold. A small number of predictions may be insufficient to capture key fold-determining inter-residues properties. However, the average number of predictions approximately corresponds to the target length with a mean error slightly greater than 1Å at a relatively high threshold probability of 0.2 (Fig. 5), suggesting an optimal configuration.

**Fig. 5.**
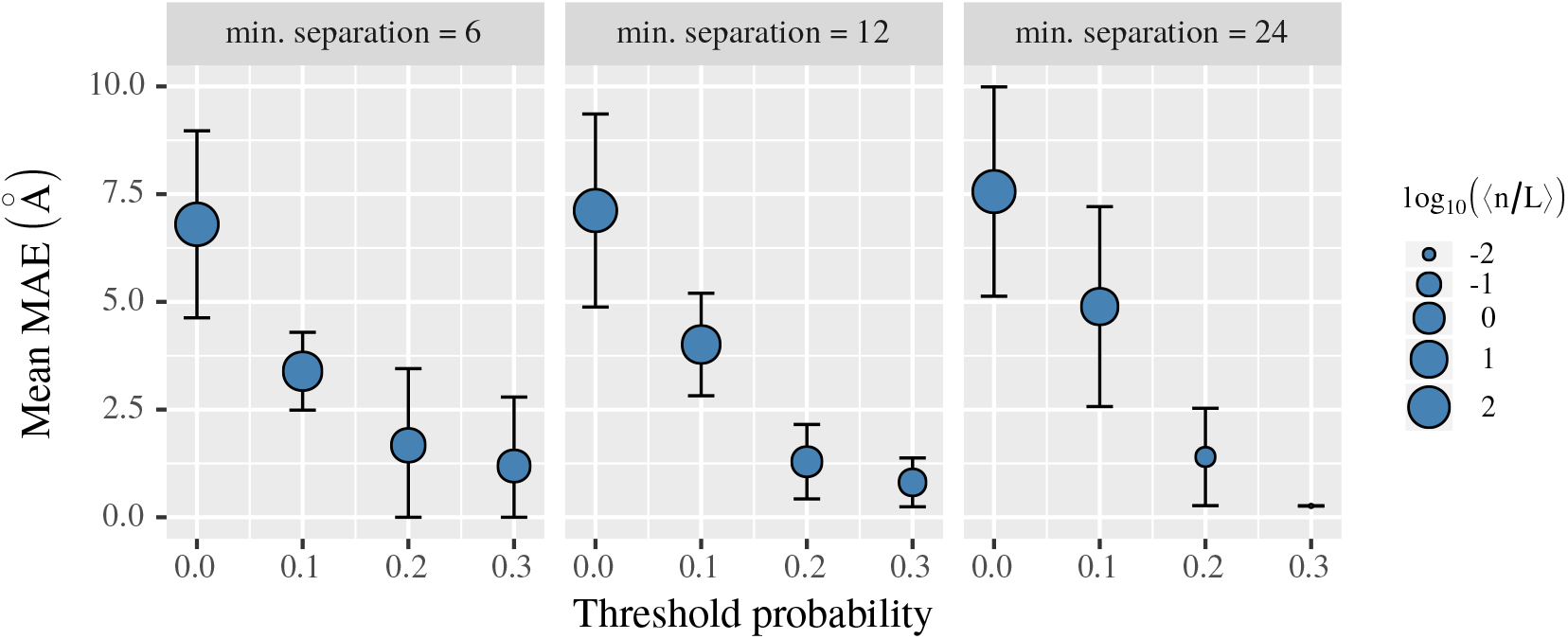
Mean of the mean absolute error (Mean MAE) between inter-residue distances predicted by the combination of the three REDCNNs and those observed in the CASP14 target structures. The mean MAE is calculated for a set of distances predicted with probability greater than the threshold probability by considering distances between any two residues separated by at least 6, 12, or 24 residues in the sequence. The point size is linearly related to the logarithm of the average number of predictions, expressed in the target length *L*, 〈*n/L*〉. (log_10_(〈*n/L*〉) = 0 implies 〈*n/L*〉 = 1.) Error bars represent the standard error of the mean MAE.

The reader is referred to Supplementary Section S1.5 (Supplementary Table S1 and S2) for a comparison of prediction accuracy with trRosetta [33].

### 3 Discussion

One of the goals of developing ROPIUS0 was to build a system for evaluating the feasibility of improving homology detection. The ROPIUS0 protocol consists of two mutually connected parts. The first is for homology modeling. The other is formed by the deep learning module designed for the selection of protein structural models. We registered ROPIUS0 as a prediction group and tested it during the CASP14 season. Its variant ROPIUS0QA aimed at unconditionally carrying out model selection was also tested. Model selection on its own is important for fold recognition, protein structure and function prediction, and modeling of protein complexes. How model selection is related to homology detection will be discussed below.

ROPIUS0QA was extensively tested and showed satisfactory results according to CASP14’s ranking in the category of tertiary structure prediction and based on additional tests on CASP13 and CAMEO targets. The architecture of the REDCNN, although being straightforward, was considered adequate for sequence-based only model selection. Its function to predict distances between residues is employed to contrast predictions with the distances observed in structural models. Differences in distances are then used to estimate the accuracy of models. Such an approach provides a reduction in computational complexity with respect to methods that use a two-level prediction system based on predicting distances first and local (residue-wise) model accuracy afterwards.

Although primarily designed for global model accuracy estimation, the REDCNN architecture can be extended to allow for multiple heads for the simultaneous prediction of distances between residues and position-specific differences in distances. Such modification would require augmenting the input format with structural models.

In general, the ROPIUS0 protocol can be transformed into an end-to-end differential framework unifying the present homology modeling and de novo modeling with model accuracy estimation. It would generate structural models from inter-residue distances and backbone dihedral angles predicted based on input data augmented with restraints transferred from identified templates (if any). Meanwhile, the discriminator component in the context of a generative adversarial network would produce model accuracy estimates for selection among the generated models.

The accuracy of modeling by homology in both a unified framework and the present study depends on the accuracy of the alignment between the target sequence and a template. ROPIUS0 applied a TSA3 algorithm (Section 5) to template selection and target-template alignment adjustment. Its effectiveness in the process of building structural models of higher quality is demonstrated in Supplementary Sections S1.1–S1.3. TSA3’s usefulness lies in general applicability, i.e., it can be applied to alignments produced by any method.

One of the main limitations of the present study arises with limited training data. While more advanced attention-based NN architectures [34] may lead to better performance, limited training data hinders learning using any NN model. This problem was overcome by focusing only on confident inter-residue distance predictions, which showed high accuracy (Figure 5). Increasing the amount of training data should lead to better model selection performance, which will be investigated in future research, but using a subset of predictions also has important implications for threading and homology search in general.

We intentionally used the COMER2 software [6] to generate sequence-based input to the REDCNN. Testing model selection using sequence-based inputs allowed us to evaluate both the accuracy of REDCNN predictions and how these predictions correlate with recognizing the correct structural fold. The result of including confident predictions offers prospects for improving the sensitivity of homology search without sacrificing much speed.

We developed a new protein threading tool COTHER and made it publicly available at https://github.com/minmarg/cother COTHER employs the COMER2 GPU-accelerated search engine but produces alignments by integrating inter-residue distance map comparison. COTHER evaluates how closely inter-residue distances predicted by ROPIUS0 for a query sequence match the distances observed in protein structures. COTHER, therefore, integrates sensitive homology detection by COMER2 and identification of common structural features predicted by ROPIUS0’s deep learning framework. This combination makes COTHER substantially more sensitive and accurate than COMER2, and the ROPIUS0 approach to focus on confident predictions makes it fast. COTHER demonstrates the application of ROPIUS0’s model selection approach to fast and sensitive sequence-based detection of evolutionary relationships between proteins, but further analysis is beyond the scope of this study.

## 4 Conclusion

In this study, we presented a protocol for protein structure prediction. Model selection represents the central part of the ROPIUS0 protocol and can be used independently to select among independently generated structural models. ROPIUS0QA, a variant based exclusively on model ranking and selection, showed satisfactory results in CASP14. The reasoning behind ROPIUS0QA is to predict interresidue distances of the protein of interest and select models whose inter-residue distances closely match confident predictions. ROPIUS0’s model selection approach was used to develop a new fast, sensitive, and accurate threading tool COTHER for sequence-based homology search and alignment. We hope the findings of this study and the ROPIUS0 and COTHER tools will be useful in protein evolution studies. ROPIUS0 and COTHER are available at https://github.com/minmarg/ropius0 and https://github.com/minmarg/cother, respectively.

## 5 Methods

This section provides details on the components of the ROPIUS0 protocol (Fig. 1).

### 5.1 Profile construction

The availability of templates is determined and templates for homology modeling are searched for by sequence profile-profile alignment. The sensitivity of the search largely depends on the quality of the profile constructed for the target sequence. To collect reliably aligned homologous sequences for a multiple sequence alignment (MSA) from which a profile is constructed, the procedure shown in Figure 6 is used.

**Fig. 6.**
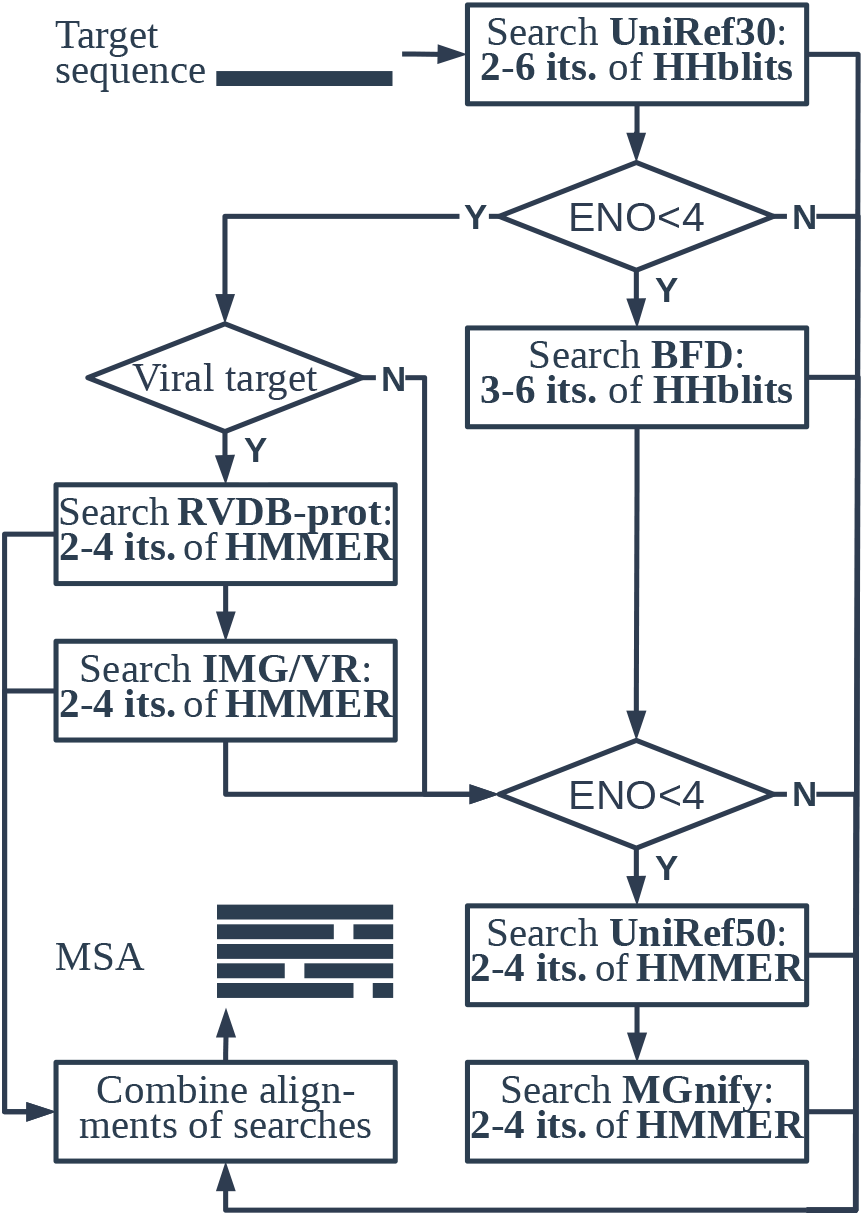
Flowchart of the procedure for generating an MSA for the target sequence.

The UniRef30 database of MSAs [28], filtered to 30% sequence identity, is searched with the target sequence using HHblits [29] for two to six iterations. If the result contains a number of sequences sufficient to construct an informative COMER2 [6] profile, no further searches are conducted. The information content of the resulting COMER2 profile is measured by the effective number of observations (ENO) [30]. Its value rarely exceeds 14, and a profile is considered to be informative if its ENO is greater than or equal to 4.

If insufficient sequences appear in the search results, HHblits is used to search the BFD metagenomic protein database of MSAs [35], filtered to 30% sequence identity. Concurrently, two viral sequence databases RVDB-prot [36] and IMG/VR [37] are searched using the jackhmmer (first iteration) and hmmsearch (subsequent iterations) programs from the HMMER3 software package [24] for two to four iterations if the target is a viral protein (Fig. 6).

If the overall search results do not imply an informative COMER2 profile, the final round of searches in the UniRef50 [38] and the MGnify metagenomic [39] sequence database is performed using the same settings for HMMER3 as described above.

All searches are configured with an *E*-value threshold for output and sequence inclusion in subsequent iterations of 0.01. The progression of iterations stops as soon as the statistical significance of a match in the previous iteration drastically decreases over subsequent iterations.

Before constructing a profile, the significant alignments from all searches conducted are combined into a single MSA. A profile is constructed from the resulting MSA using the makepro program from the COMER2 software package.

### 5.2 Template search

For identifying structural templates, the PDB70 database [40] of COMER2 profiles corresponding to PDB [41] sequences filtered to 70% identity is searched using COMER2 [6] with a query profile constructed for the target sequence. The output of the search is controlled by the option MINPP of a posterior probability threshold for alignment extension. MINPP values of 0.28 (default) and 0.1 were used, and alignments produced using a value of 0.1 were selected if the aligned query coverage was not much larger than that obtained using the default value. A MINPP value of 0 was not used, as semi-global alignments produced in this setting can include non-homologous, structurally divergent terminal regions.

### 5.3 Template selection and alignment adjustment algorithm TSA3

An algorithm for template selection and alignment adjustment TSA3 has been developed (Fig. 7). The rationale behind the algorithm is that statistically significantly identified templates with the most extensive alignment coverage considered reliable represent the most appropriate candidates for protein homology modeling. The regions considered reliably aligned between a template and the target result from applying the TSA3 algorithm to that template. The algorithm applies to every template of interest individually.

**Fig. 7.**
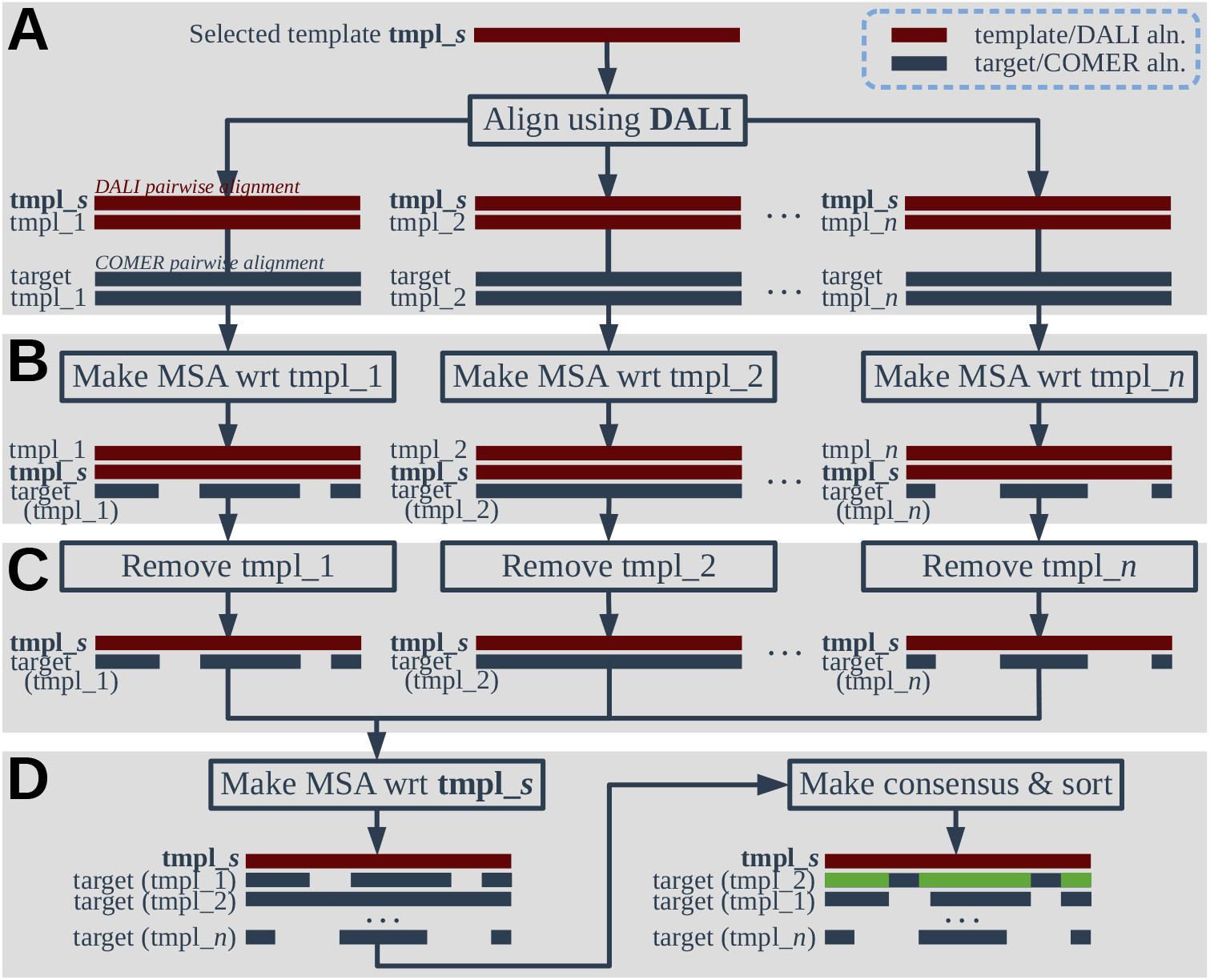
TSA3 algorithm for template selection and alignment adjustment. Its goal is to identify reliably aligned regions for each template tmpl_*s* of interest. The algorithm starts with producing the pairwise structural alignments between tmpl_*s* and all members of the template set {tmpl_*i*, using DALI (A). Next, for each template tmpl_*i* (*i* = 1,...,*n*), the MSA is derived from the DALI pairwise alignment between tmpl_*s* and tmpl_*i* and the COMER pairwise alignment between the target and tmpl_*i* (target (tmpl_*i*); B). Template tmpl_*i* is removed from the MSA to leave tmpl_*s* and the target associated through it in the pairwise alignment (C). Consensus analysis of the target alignment variants in the MSA derived from the indirect pairwise alignments between tmpl_*s* and the target reveals reliably aligned regions (green; D). The TSA3 algorithm applied independently to multiple templates tmpl_*s* suggests for protein modeling candidates with extensive reliably aligned regions.

Let tmpl_*s* denote the template whose alignment with the target sequence is to be adjusted by applying the TSA3 algorithm. Let *T* be the set of *transition* templates tmpl_*i* homologous to both tmpl_*s* and the target. According to the algorithm, reliably aligned regions are aligned consistently across the alignment variants between tmpl_*s* and the target. The tmpl_*s*-target alignment variants are obtained by first aligning the transition templates tmpl_*i* with both the target and tmpl_*s*. For each tmpl_*i* ∈ *T*, two pairwise alignments, tmpl_*i*-tmpl_*s* and tmpl_*i*-target alignments, are merged to form the MSA using tmpl_*i* as a reference sequence. Leaving tmpl_*i* out then produces one tmpl_*s*-target alignment variant (Fig. 7).

The accuracy of the TSA3 algorithm depends on pairwise alignment quality. Since one set of pairwise alignments includes template sequences for which the structure is known, structure comparison, considered the most accurate, is used to produce the alignments between tmpl_*s* and all the templates tmpl_*i* ∈ *T*. We use DALI [42].

A schematic of the TSA3 algorithm is shown in Figure 7. In addition to indicating reliably aligned regions, alignment variants sorted by proximity to the consensus alignment offer alternatives for adjusting the alignment between the target and tmpl_*s* to be used for homology modeling. The TSA3 algorithm is most helpful when the set *T* includes structurally similar templates.

The approach of using structural alignments to improve alignment accuracy is not new [43]. The main differences of the TSA3 algorithm are that the alignment between the target and the template of interest is adjusted based on their dependent association (multiple MSAs), and reliably aligned regions remain unchanged (no realignment). The algorithm is readily applied to one template.

### 5.4 Homology modeling

Homology modeling is used to predict the structure of the target sequence when structural templates are available (Section 5.2). Modeling is performed with RosettaCM [44] version 2019.35.60890 using the ref2015 score function [45], which is applied to modeling every target domain using optionally multiple templates and to combining domain structural models into a complete structure of the target. Target domain boundaries are identified using COMER2 and from REDCNN predictions (see Section 5.5). One or more templates for modeling are selected and their alignments to the target are adjusted using the TSA3 algorithm (Section 5.3) if multiple templates are available. Five to eight selected structural models (method described in Section 5.8 and used in the second half of the CASP14 season) of each identified target domain are refined using the Rosetta relax [46] computer program. Relaxed models were not considered if they were more divergent from the template(s) than the selected starting models. The number of decoys generated by RosettaCM and relax was 200. The same refinement using relax is applied to selected CASP-hosted server models when templates are unavailable.

### 5.5 REDCNN

A REDCNN is used to predict a probability distribution of distances between the CB atoms of each pair of the target residues. Its fully convolutional architecture, shown in Figure 8, is based on SegNet [47]. It consists of the encoder and decoder parts. The encoder contains a series of groups of blocks, each followed by a maxpool layer. The max-pooling operation applied with a window of size 2 × 2 and a stride of the same size halves the dimensions of each two-dimensional (2D) input feature map. Each block in a group preceding a maxpool layer contains a convolution layer followed by batch normalization and ReLU activation.

**Fig. 8.**
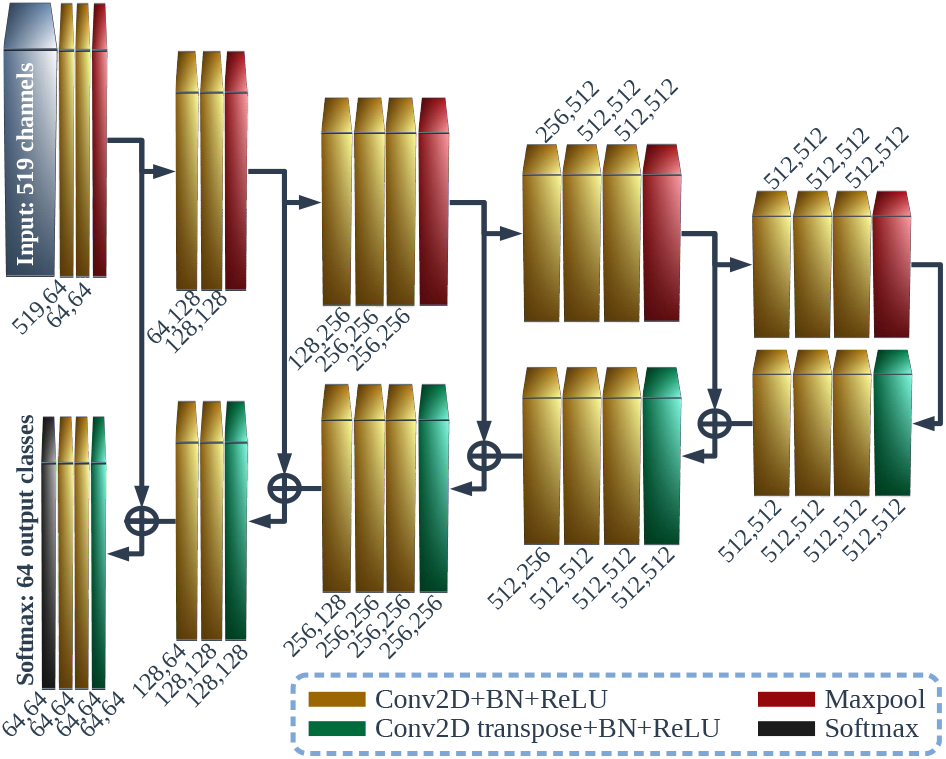
REDCNN architecture. Two numbers below or above blocks indicate the number of input feature maps (channels in the first layer) and the number of filters in the convolution and the convolution transpose layers, respectively. For the softmax layer, the second number is the number of classes. Kernels of size 3 × 3 are used for convolution and transposed convolution. Input has 519 channels. Output depth corresponds to a probability distribution over 64 classes predicted by the last softmax layer. BN, batch normalization.

The decoder has an upscaling structure. In the decoder, a convolution transpose layer, batch normalization and ReLU activation precede each group of blocks organized similarly as in the encoder. A stride of size 2 × 2 used for transposed convolution ensures the dimensions of an output feature map agree with those of a feature map fed into the maxpool layer at the corresponding level in the encoder. The units of each feature map produced by a maxpool layer of the encoder are connected by skip connections to the units of the corresponding feature map passed to a convolution transpose layer of the decoder (Fig. 8).

A probability distribution of distances is predicted by the softmax layer. The output for a length-L target protein or its fragment is an *L × L* map whereby each cell is a probability distribution over 64 classes, which represent distances discretized into intervals of length 1A ([0, 2), [2, 3),..., [63, 64), [64, ∞)). Clusters of relatively short (mean) distances in local regions of the predicted map can provide additional indication of domain boundaries for the target protein.

### 5.6 REDCNN input

Multiple *L × L* attribute matrices stacked one on top of another constitute the input to a REDCNN. Each matrix represents a specific type of data for a target or its fragment of length *L*. A matrix entry indexed by *i* and *j* contains a value that characterizes a relation between residues at positions *i* and *j*. The number of matrices is called the number of channels in the input.

There are two groups of attribute matrices. The first group is obtained from a COMER profile constructed from an MSA representing the family of the target sequence (Section 5.1). Profile data extends along the target sequence.

Therefore, the value of a profile attribute at position *i* is assigned to row *i* and replicated across all columns of an attribute matrix. Accordingly, the same value at profile position *i* is additionally associated with column *i* and replicated across all rows to obtain a 2D representation of profile data.

Profile attributes at position *i* and the number of channels required are the following: 1 channel of sequence position index; 1 channel of residue code; 20 channels of amino acid substitution probabilities (probability distribution ***f**_i_*) [48]; 7 channels of transition probabilities from and to match (M), insert (I), and delete (D) states; 3 channels of effective numbers of observations; 1 channel of the sum of the five largest posterior cluster membership probabilities and the probability of a new cluster for ***f**_i_* under the hierarchical Dirichlet process (HDP) mixture model [48]; 2 channels of the largest cluster membership probability under the HDP mixture model and the index of the corresponding cluster; 2 channels of the squared Euclidean norm and the logarithm of the prior probability of the context vector [49]; 19 channels of the context vector itself; 3 channels of secondary structure probabilities for coil (C), strand (E), and helix (H) classes [50].

The second group of attribute matrices is 2D by nature and corresponds to the mutual information and the cross-covariance matrix between observations at positions *i* and *j* of the MSA (and target sequence) for each *i* and *j*.

The cross-covariance matrix of two random vectors ***v**_i_* and ***v**_j_* of dimension 20 is

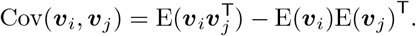

We estimate it from a sample of sequences, as found in the MSA, that have residues aligned at positions *i* and *j*. An observation ***v**_sk_* of random variable ***v**_k_* records an occurrence of the residue found in sequence *s* at position *k*. The ath element of ***v**_sk_* is *v_ska_* = *n_k_w_s_*, where *n_k_* is the number of sequences at position *k* in the MSA, and *w_s_* (Σ_*s*_ *w_s_* = 1) is the weight of sequence *s* with residue type *a* at position *k* [51]. The total number of channels for the cross-covariance matrix is 400 (20 × 20).

Mutual information is calculated as the Kullback-Leibler divergence between estimated probability distributions *D*_KL_(*p_ij_*|| *q_ij_*), where the joint probabilities of observing residue types *a* and *b* at positions *i* and *j* are

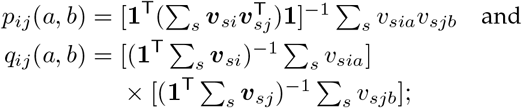

**1** is a vector of ones. Mutual information requires one channel.

With a total of 519 channels, input to a REDCNN is produced using the COMER2 software package. The makepro program constructs a profile from an MSA, whereas the makecov program calculates cross-covariance matrices and mutual information for all pairs of positions *i* and *j* (*i* ≤ *j*). We refer to this type of input as a *promage*.

### 5.7 Combining multiple REDCNN predictions

We apply three trained instances of the REDCNN model (Supplementary Section S2.1, Supplementary Fig. S8) to predicting inter-residue distances and model accuracy estimation (Section 5.8). In the former case, predictions made by three REDCNNs are combined to form a mixture of probability distributions with equal weights for each pair of residues.

### 5.8 REDCNN-based model accuracy estimation

A trained REDCNN model is used to estimate the accuracy of a protein structural model of length *L.* Accuracy estimates are produced by evaluating how close the distances between CB atoms in the structural model match those predicted by the REDCNN.

Residue-wise accuracy estimates are calculated for each protein residue and represent errors in Angstroms, as required by CASP. An estimate for residue *i* (*i* ∈ [1,*L*]) is obtained by first calculating the mean absolute difference between distances observed in the structure, and rounded to the nearest integer, and predicted distances over pairs of residues (*i, j*) (*j* = 1,..., *i*—4, *i*+4,..., *L* for 4 < *i* ≤ *L* — 4) such that they are predicted, with probability at least the threshold probability, to be separated by a distance at most the threshold distance. The mean absolute difference obtained for residue *i* is divided by the expected difference for random structures (= 1/63 × 63 · 62/2) and multiplied by 8 so that the expected error is 8A. We additionally multiply the estimate by 2 to account for using the thresholds and for a relatively small probability of long distances between residues in protein structures.

The global accuracy score provides an estimate for the entire protein structure. It is calculated as 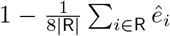, where 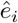 is the accuracy estimate calculated for residue *i*, and R corresponds to the set of residues included in the calculation, whose size is not necessarily *L* due to the thresholding and the possibility of missing residues in the structural model. The global accuracy estimate ranges from 0 to 1.

The calculation of final global and local estimates is based on averaging the estimates obtained using three trained instances of the REDCNN model (Supplementary Section S2.1). The estimates also depend on the physics of the structural model evaluated using Rosetta [46]. The final local estimate for residue *i* is, therefore, a linear combination of the average estimate and the exponentiated sum of certain Rosetta energy terms for residue *i*, with weights 1 — *r* and *r* (0 < *r* < 1), respectively. The final global estimate is calculated as described above, using the final local estimates.

The set of parameters that applies to calculating the accuracy estimates includes the probability and distance thresholds for each of the three REDCNNs, the list of Rosetta energy terms, and the weight *r*. We determined their optimal configuration on a small subset of CASP13 targets.

We distinguished between two conditions/modes. These conditions represent the situations of abundance and scarcity of sequences homologous to the sequence of the protein under consideration. They also represent the CASP template-based (TBM) and free modeling (FM) categories for target sequences.

Two groups of five targets served as datasets. The first group of targets for which numerous homologous sequences could be found consisted of the following targets: T0965, T0976, T0996, T0997, and T1014. The targets in the second group were T0950, T0987, T0990, T1005, and T1010. They represented sequences of the FM category.

The optimization criterion was set to achieve the best match between the rankings of the top 50 structural models ranked officially by CASP13 according to GDT_TS [20] and the rankings obtained by the final global estimate for these structural models. The weight *r* = 0.2 and a list of four Rosetta energy terms, fa_elec, hbond_lr_bb, hbond_sc, and p_aa_pp, were found to be optimal for both groups of targets. The probability and distance threshold values for each of the three REDCNNs were found to be (0.2, 32), (0.2, 32), and (0.2, 20) for the first group of targets. The corresponding values for the second group of targets were (0.3, 20), (0.1,10), and (0.1, 8).

## Acknowledgments

The author is grateful to Albertas Timinskas for help in preparing data for the experiments.

## Software and Data Availability

The ROPIUS0 software is available at https://github.com/minmarg/ropius0 Trained TensorFlow models are deposited to Zenodo at https://doi.org/10.5281/zenodo.4450107 Code, scripts, and data for tests on CASP datasets are deposited to Zenodo at https://doi.org/10.5281/zenodo.4451302 and https://doi.org/10.5281/zenodo.4783276 CASP and CAMEO datasets are available at https://predictioncenter.org and https://www.cameo3d.org, respectively.

## S1 Supplementary Results

### S1.1 TSA3 application in CASP14

The TSA3 algorithm for template selection and alignment adjustment as part of the ROPIUS0 protocol was applied during the CASP14 season. To investigate how effective the TSA3 algorithm is for homology modeling, additional experiments were conducted after the CASP14 season had ended.

As COMER2 alignments had been adjusted by the TSA3 algorithm, the unadjusted COMER2 alignments were used for evaluation. The most accurate model, among the five submitted models by the ropius0 group, was compared with the most accurate model obtained from modeling using the five most significant templates identified by COMER2 (Margelevičius, 2020) for each CASP14 target for which the structure is available and TSA3 had been applied during the CASP14 season. (TSA3 applies to query sequences for which multiple homologous templates can be identified.) Modeling of full targets was performed using RosettaCM (Song *et al*., 2013) (see the section “Homology modeling” of the main text). For each of the identified templates, 20 decoys were generated. The most accurate model among the models generated using each of the five templates was considered the model obtained using the COMER2 alignment for an individual target. The sequence-dependent superposition between a model and the target, and their structural similarity by GDT_TS score, was calculated using LGA Zemla (2003) with the same parameters as used by CASP (Protein Structure Prediction Center, 2020). The evaluation was performed on a full-target and target domain basis.

The results are shown in Figure S1. We note that model selection was based on Rosetta energy scores during the execution of the ROPIUS0 protocol in the first half of the CASP14 season. The deep learning-based model selection had been applied to one target, T1099, from the set of targets used in this analysis. The domains of the multi-domain targets of this set had not been modeled separately.

Figure S1 shows that the application of the TSA3 algorithm leads to an increase in the average GDT_TS score. The increase, though, is insignificant due to the small sample size (*p*-value = 0.065, one-sided Wilcoxon signed-rank test).

### S1.2 TSA3 application to CASP13 targets

An additional experiment was performed to directly evaluate the effect of the TSA3 algorithm on modeling CASP13 targets in the TBM (template-based modeling) category. A target was modeled in two ways. According to the first, a structural model was generated using a COMER2 alignment between the target sequence and a template. The second way of modeling was based on the COMER2 alignment between the target and the same template, adjusted using the TSA3 algorithm. The accuracy of obtained models for the same target and using the same template allowed us to evaluate TSA3’s impact on alignment accuracy.

Templates were searched using COMER2 as specified in the section “Template search” of the main text. Templates identified for a target sequence were not considered if they had been released after the target sequence release during the CASP13 season (2018). Among the top-scoring candidate templates identified by COMER2, one with the highest sequence identity to the target sequence was selected as a template for modeling. If the selected template did not lead to adjustments in the alignment, the next best template was selected. RosettaCM was used to generate 20 full-length models for each target. The models were evaluated on a target domain basis using the LGA sequence-dependent superposition between the target and the model with the highest Rosetta score among the generated decoys. Long insertions (>4 residues) in the models were removed so that free modeling did not affect model evaluation.

**Figure S1.**
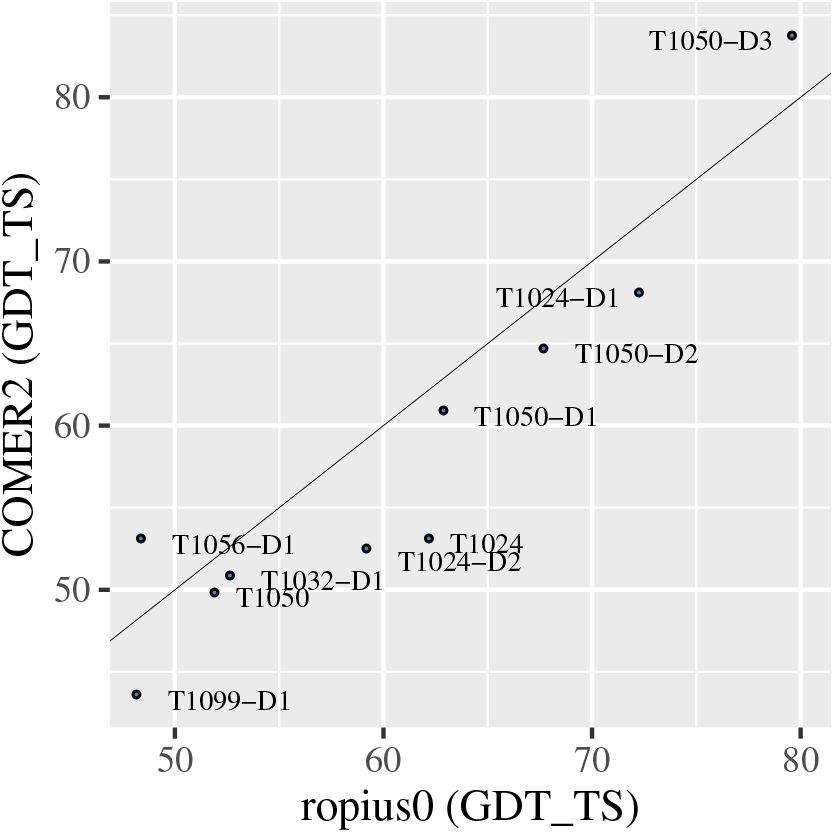
Accuracy of the structural models obtained solely using COMER2 alignments and the accuracy of the models submitted by the ropius0 group for a subset of the available CASP14 target structures. Model accuracy is evaluated by the GDT_TS score. The straight line represents the equivalent model quality.

Figure S2 shows the results for the targets for which adjustments suggested by the TSA3 algorithm applied. Consistent improvement in the GDT_TS score is observed for all but one domain, T0961-D1: 87.58 vs. 87.53. The improvement is statistically significant (p-value = 3.1 × 10^-5^, one-sided Wilcoxon signed-rank test). This test demonstrates the effectiveness of the TSA3 algorithm, which is not limited to COMER2 alignments and can be applied to alignments produced by any method.

As an example, the results of applying TSA3 to the HHsearch alignments (Steinegger *et al*., 2019) using the same evaluation process for the CASP13 targets are shown in Figure S3. A statistically significant improvement is observed (p-value = 5.0 × 10^-4^).

### S1.3 Examples of TSA3 application

This section provides two examples illustrating the benefit of the TSA3 algorithm. The first example is the modeling of target T1016 using PDB (Berman *et al*., 2000) template 1bif_A. Consensus analysis by the TSA3 algorithm suggested an insertion of two residues, with respect to the original alignment, in a beta strand at the C-terminus (Figure S4A). The adjustment corrected the alignment. Using the corrected alignment resulted in a more accurate structural model with an improvement in GDT_TS of 7.3.

Figure S4B shows the second example, the result of modeling target T1005 using PDB template 4nuz_A. This time, TSA3 highlighted a reliably aligned alpha helix at the C-terminus, which was missing in the original alignment. This region was not evaluated in the analysis of Section S1.2 because it represented a long insertion. The improvement in GDT_TS was, therefore, 0.9. Model accuracy improved by 4.5 GDT_TS points when the original alignment was extended by appending the aligned region suggested by the TSA3 algorithm (Figure S4B).

**Figure S2.**
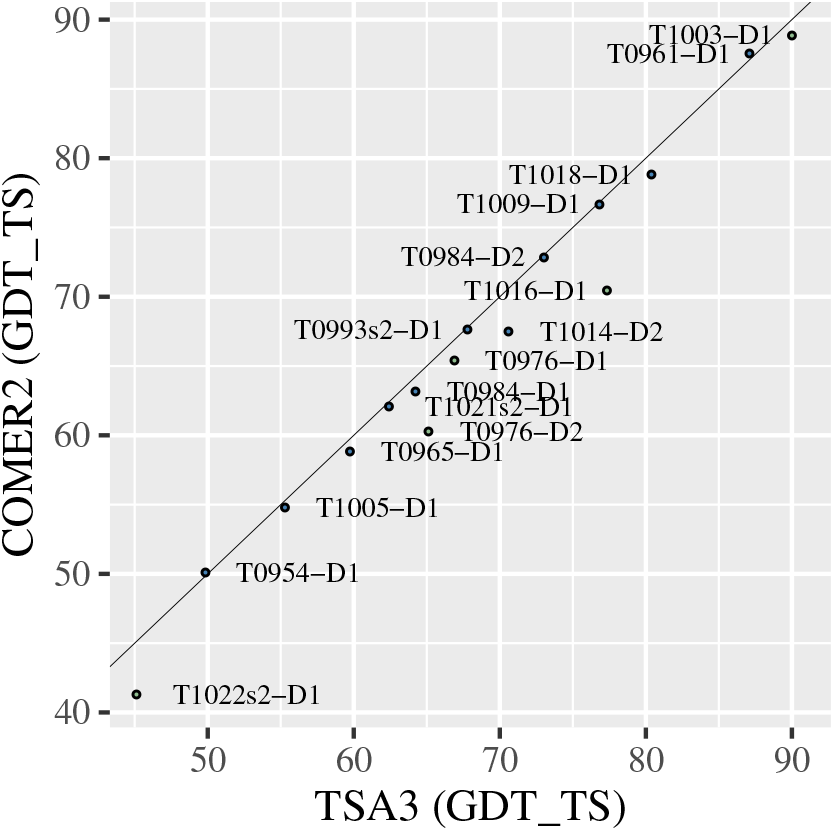
Accuracy of the structural models obtained using COMER2 alignments and alignments adjusted by the TSA3 algorithm for CASP13 targets in the TBM category. The green points denote models obtained not using templates selected first. Model accuracy is evaluated by the GDT_TS score. The straight line represents the equivalent model quality.

**Figure S3.**
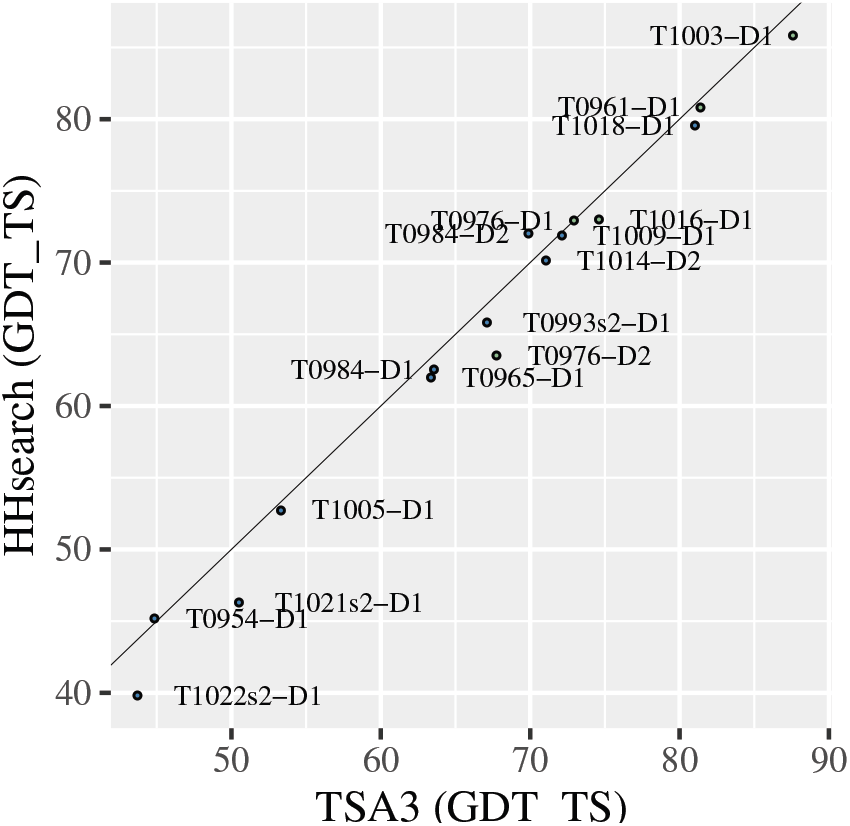
Accuracy of the structural models obtained using HHsearch alignments and TSA3-adjusted alignments for CASP13 targets in the TBM category. The green points denote models obtained not using templates selected first.

**Figure S4.**
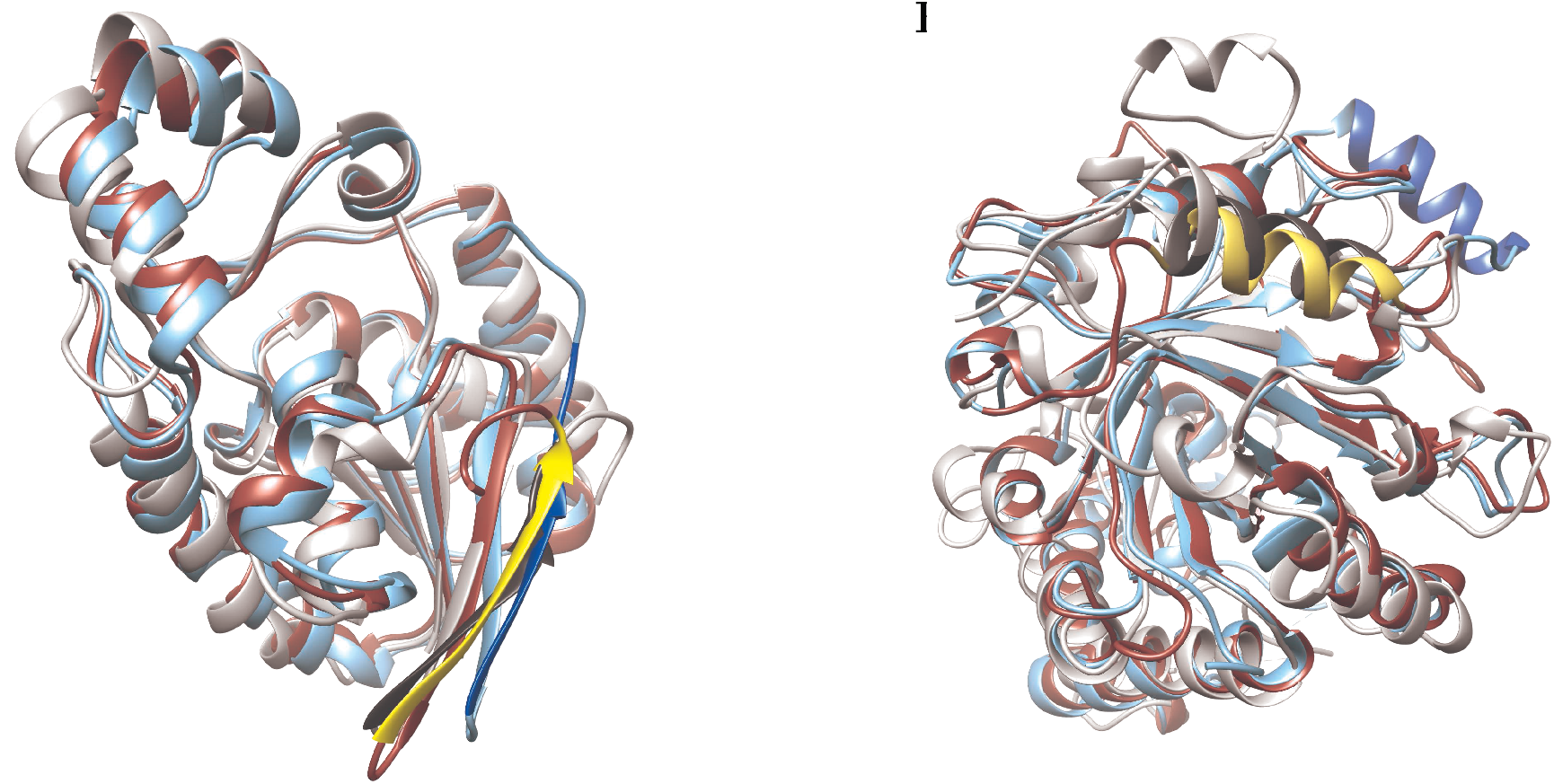
Modeling examples. (A) Modeling CASP13 target T1016 using template 1bif_A. (B) Modeling T1005 using template 4nuz_A. The target structure is colored light grey. The model obtained using the COMER2 alignment is in light blue. The model obtained using the alignment adjusted by TSA3 is displayed in red. The matching regions discussed in the text are highlighted in a darker/brighter shade of the corresponding color. The region of the target is in dark grey. Yellow highlights accuracy improvement from the adjustments. Images prepared using UCSF Chimera (Pettersen *et al*., 2004).

### S1.4 ROPIUS0QA’s performance on CASP13 targets

ROPIUS0QA was tested on the CASP13 targets. Three other methods DeepAccNet (Hiranuma *et al*., 2021), ProQ3D (Uziela *et al*., 2017), and QMEANDisCo (Studer *et al*., 2020) were evaluated too.

The results for DeepAccNet were obtained from the available predictions on the CASP13 dataset (Hiranuma *et al*., 2021). ProQ3D was provided with the UniRef90 database (Suzek *et al.*, 2015) dated January 1, 2021. The setup for QMEANDisCo was as follows. QMEANDisCo uses template structures and the alignments between the target sequence and the templates for estimating model accuracy. Templates were identified by searching the HHsuite PDB70 database dated March 10, 2021, using HHsearch (Steinegger *et al*., 2019). The query (target) profile HMMs were constructed from the same multiple sequence alignments (MSAs) with which ROPIUS0QA was provided. The secondary structure for the target was predicted using PSIPRED (Jones, 1999). Solvent accessibility was predicted using ACCpro (SCRATCH-1D) (Magnan and Baldi, 2014). The number of top templates identified with a significance threshold of E-value = 0.01 was limited to 5. Each template sequence was aligned with the structure to ensure the consistent alignment between the target and template sequences.

The ROPIUS0QA results are shown in Figure S5. The evaluation results are also reported in Table 2 (main text), which includes the results of the ROPIUS0QA versions that used the Uniclust30 database (Mirdita *et al.*, 2017) released before the start of the CASP13 season and the then version (3.0.0) of HHblits (Remmert *et al*., 2012). The runtimes of the evaluated methods for all targets are shown in Figure S6.

**Figure S5.**
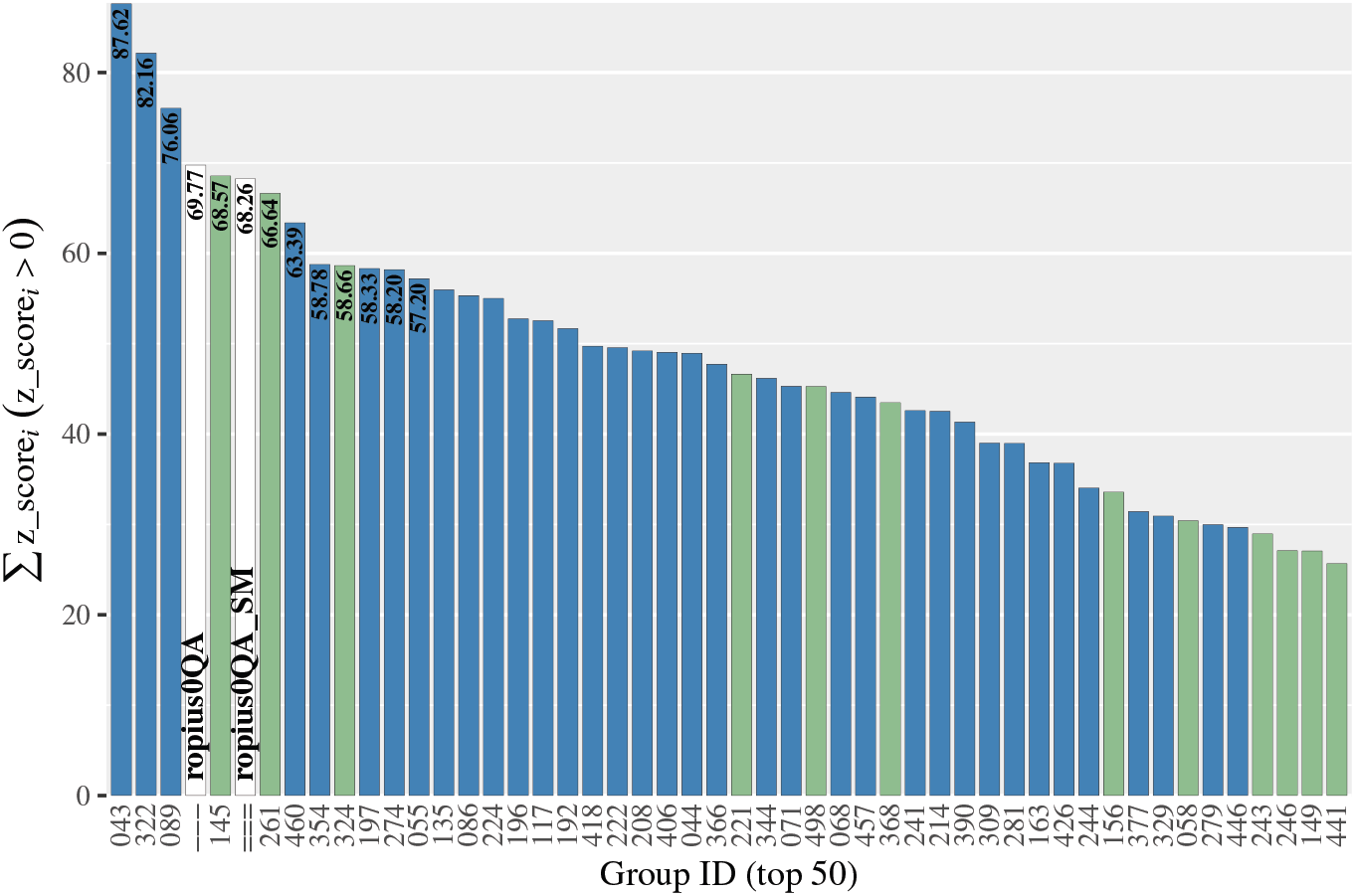
Results of testing ROPIUS0QA on CASP13 targets (76 domains) in the tertiary structure prediction category. Ten targets used to determine thresholds for the REDCNNs are excluded. Five more targets/domains (T0954, T0966, T0983, T0999, and T1017s1-D1) having sequence similarity to a member of the training set are excluded too. The ranking is based on the sum of the *z*-scores found to be positive for the structural models designated by the group under consideration as best (1st model). ropius0QA_SM stands for single-mode ROPIUS0QA (see Methods in the main text). The top 50 groups are shown. Server prediction groups are represented in green.

**Figure S6.**
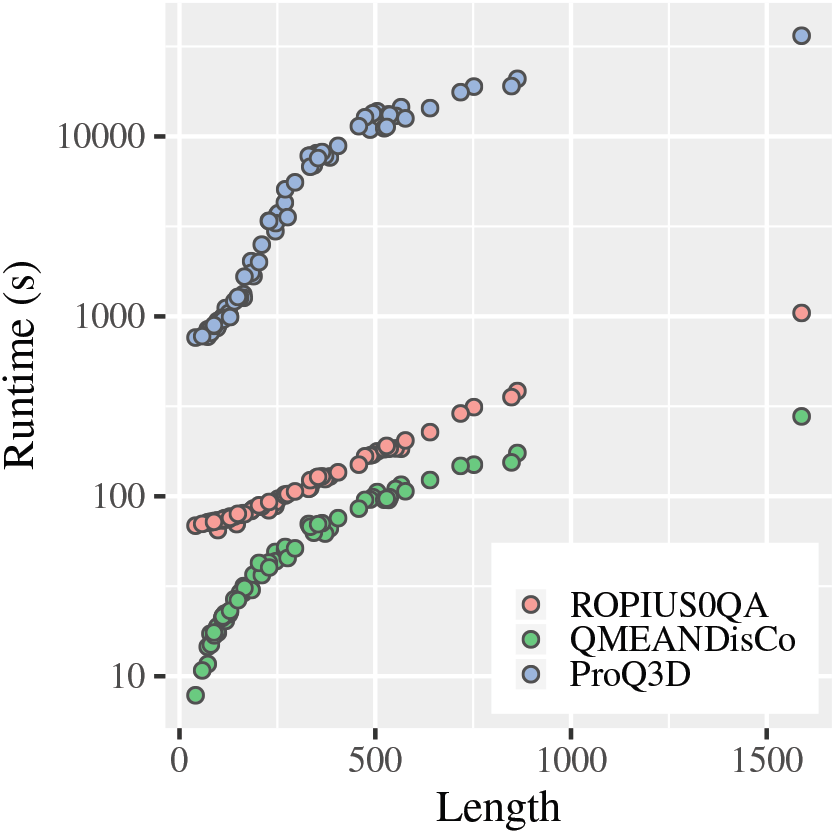
Runtime for all CASP13 targets (104 domains), excluding the time spent on sequence search. The time spent on the template search, target-template alignment, and external predictions is not included for QMEANDisCo.

### S1.5 REDCNN prediction accuracy

**Figure S7.**
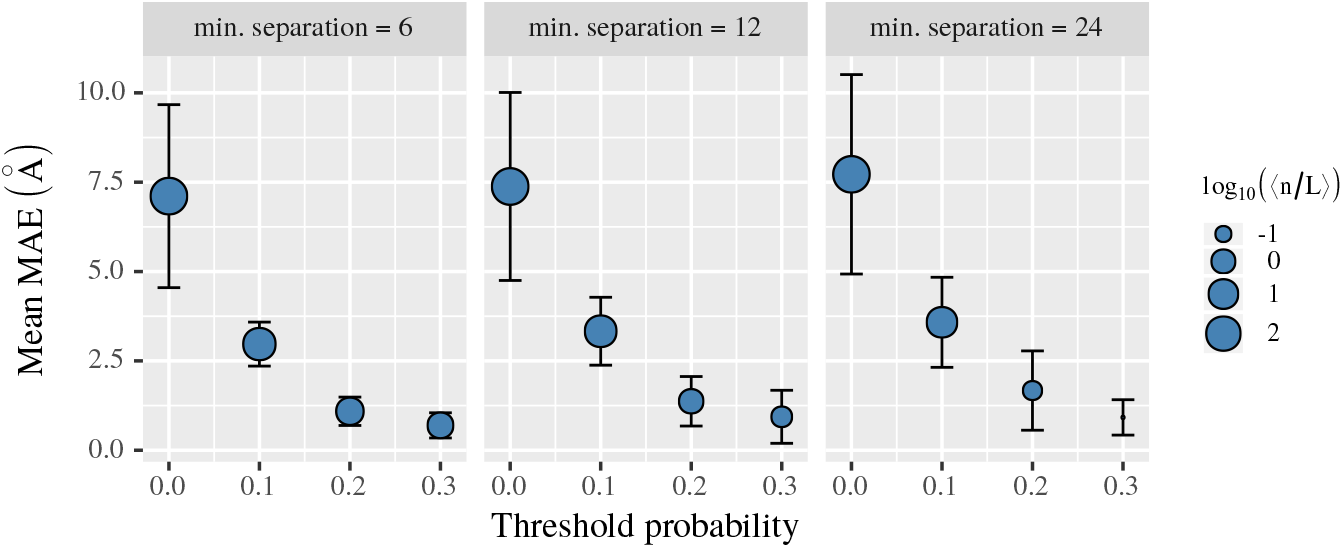
Mean of the mean absolute error (Mean MAE) between inter-residue distances predicted by the combination of the three REDCNNs and those observed in the CASP13 target structures. The mean MAE is calculated for a set of distances predicted with probability greater than the threshold probability by considering distances between any two residues separated by at least 6, 12, or 24 residues in the sequence. The point size is linearly related to the logarithm of the average size of a set of distances expressed in the target length *L*, {*n/L*}. Error bars represent the standard error of the mean MAE.

**Table S1.** Mean of the mean absolute error (Mean MAE, Å) of inter-residue distances predicted by ROPIUS0/ROPIUS0QA (by the combination of the three REDCNNs) and trRosetta for the CASP14 targets. For each target, the same MSA is given as input to both methods. The mean MAE is calculated for a set of distances predicted with probability greater than the threshold probability (Prb) by considering distances between any two residues separated by at least 6, 12, or 24 residues in the sequence (min. sep.). The mean number of predictions, expressed in the target length L, is denoted by 〈*n/L*〉. s.e. is standard error. trRosetta was trained on 15,051 protein structures (Yang *et al*., 2020); its resolution of prediction is 0.5^Å^. Note a difference in the number of predicted distances (〈*n/L*〉) between the methods. The number of the trRosetta predictions is more uniformly distributed at low probability values. The number of the ROPIUS0 predictions with a probability just above 0.0 is almost *L*^2^ (〈*n/L*〉 ≈ averge_protein_length) and can be considered noise. Therefore, predictions of low confidence are ignored (see the main text).

|  |  | ROPIUS0 |  | trRosetta |  |
| --- | --- | --- | --- | --- | --- |
| | | Mean MAE (s.e.) | $\langle n/L \rangle$ (s.e.) | Mean MAE (s.e.) | $\langle n/L \rangle$ (s.e.) |
| min. sep.=6 | Prb=0.0 | 6.80 (2.17) | 254.43 (445.51) | 2.33 (1.23) | 24.32 (32.64) |
|  | Prb=0.1 | 3.39 (0.90) | 31.24 (55.94) | 1.81 (1.13) | 14.82 (19.82) |
|  | Prb=0.2 | 1.67 (1.78) | 3.53 (4.76) | 1.29 (1.08) | 4.40 (5.03) |
|  | Prb=0.3 | 1.19 (1.60) | 1.89 (2.71) | 0.96 (1.03) | 2.49 (2.67) |
| min. sep.=12 | Prb=0.0 | 7.12 (2.24) | 242.54 (436.89) | 2.66 (1.97) | 17.03 (27.71) |
|  | Prb=0.1 | 4.01 (1.19) | 21.64 (48.60) | 2.29 (2.02) | 9.54 (16.00) |
|  | Prb=0.2 | 1.29 (0.86) | 0.86 (1.56) | 1.30 (0.99) | 1.40 (2.39) |
|  | Prb=0.3 | 0.81 (0.57) | 0.28 (0.53) | 0.81 (0.45) | 0.50 (0.77) |
| min. sep.=24 | Prb=0.0 | 7.56 (2.43) | 219.86 (419.72) | 2.87 (3.31) | 11.64 (19.67) |
|  | Prb=0.1 | 4.89 (2.32) | 11.79 (37.40) | 2.55 (3.50) | 6.24 (10.73) |
|  | Prb=0.2 | 1.40 (1.13) | 0.04 (0.18) | 1.05 (0.76) | 0.54 (0.83) |
|  | Prb=0.3 | 0.26 (0.00) | 0.01 (0.04) | 1.01 (0.96) | 0.14 (0.24) |

**Table S2.** Mean of the mean absolute error (Mean MAE, Å) of inter-residue distances predicted by ROPIUS0/ROPIUS0QA and trRosetta for the CASP13 targets. See Table S1 for details.

|  |  | ROPIUS0 |  | trRosetta |  |
| --- | --- | --- | --- | --- | --- |
| | | Mean MAE (s.e.) | $\langle n/L \rangle$ (s.e.) | Mean MAE (s.e.) | $\langle n/L \rangle$ (s.e.) |
| min. sep.=6 | Prb=0.0 | 7.11 (2.56) | 370.44 (625.33) | 1.80 (0.92) | 28.41 (21.70) |
|  | Prb=0.1 | 2.97 (0.62) | 36.31 (42.62) | 1.40 (0.64) | 18.49 (14.13) |
|  | Prb=0.2 | 1.09 (0.40) | 3.80 (4.93) | 0.91 (0.40) | 4.61 (3.17) |
|  | Prb=0.3 | 0.70 (0.36) | 1.98 (3.03) | 0.73 (0.33) | 2.43 (1.93) |
| min. sep.=12 | Prb=0.0 | 7.38 (2.63) | 357.27 (614.61) | 2.00 (2.02) | 21.41 (17.45) |
|  | Prb=0.1 | 3.33 (0.95) | 26.14 (34.14) | 1.49 (1.01) | 13.43 (11.11) |
|  | Prb=0.2 | 1.37 (0.69) | 1.18 (1.73) | 0.95 (0.46) | 2.05 (1.53) |
|  | Prb=0.3 | 0.94 (0.74) | 0.33 (0.65) | 0.76 (0.39) | 0.76 (0.56) |
| min. sep.=24 | Prb=0.0 | 7.72 (2.79) | 331.84 (593.28) | 2.41 (4.43) | 15.80 (14.20) |
|  | Prb=0.1 | 3.58 (1.26) | 15.07 (22.50) | 1.98 (4.04) | 9.65 (8.95) |
|  | Prb=0.2 | 1.67 (1.11) | 0.21 (0.40) | 0.91 (0.57) | 1.20 (1.09) |
|  | Prb=0.3 | 0.92 (0.50) | 0.02 (0.06) | 0.75 (0.68) | 0.39 (0.39) |

### S1.6 Benchmarking on CAMEO targets

ROPIUS0QA was also benchmarked on a dataset of 186 CAMEO (Robin *et al*., 2021) targets released over three months, from 07/24/2021 through 10/16/2021, in the category of structure prediction. MSAs from which the input to the REDCNNs was generated were obtained by searching the UniRef30 2021_06 database (Mirdita *et al*., 2017) with the target sequences using HH-blits. The exceptions were targets 2021-07-31_00000268_1 (7RJU_A) and 2021-08-28_00000135_1 (7FE0_A), where we additionally used HMMER3 (Eddy, 2011) to search the UniRef50 2021_03 sequence database (Suzek *et al*., 2015), and target 2021-09-18_00000068_1 (7JTA_A), which was searched against the MGnify 2019_06 metagenomic sequence database (Mitchell *et al*., 2020) using HMMER3. ROPIUS0QA was then applied to the first structural models (designated by the servers as best) submitted by the registered prediction servers for each target. The ROPIUS0QA selection was the model ranked the highest.

The results of the single-mode version of ROPIUS0QA (both versions performed similarly) are shown in Table S3. The results show that ROPIUS0QA’s deep learning framework performs well in selecting accurate models. In general, the results of the three different tests—blind testing in CASP14 (main text), benchmarking on CASP13 and CAMEO datasets—consistently show good model selection performance.

**Table S3.** Benchmark results on the CAMEO dataset. ROPIUS0QA (boldface) selected among the servers’ models. #Modeled, the number of targets modeled by the server; lDDT_all, the average lDDT score (Mariani *et al*., 2013) calculated over all released (186), hard (22), or medium (99) targets; lDDT_mod, the average lDDT score over modeled targets; TM_all, the average TM-score (Zhang and Skolnick, 2004) over all released, hard, or medium targets; TM_mod, the average TM-score over modeled targets. Hard targets are those for which the average lDDT score of the corresponding structural models (ROPIUS0QA excluded) is less than 0.50. The average lDDT score of medium targets falls in the interval [0.5, 0.75). The methods are sorted by lDDT_all obtained for all released targets. ^1^BestSingleStructuralTemplate; ^2^Naive AlphaFold-EBI 90; ^3^Naive AlphaFold-EBI 100.

| Server | ALL TARGETS |  |  |  |  | HARD |  |  | MEDIUM |  |  |
| --- | --- | --- | --- | --- | --- | --- | --- | --- | --- | --- | --- |
|  | #Modeled | IDDT_all | IDDT_mod | TM_all | TM_mod | #Modeled | IDDT_all | IDDT_mod | #Modeled | IDDT_all | IDDT_mod |
| RoseTTAFold | 186 | 0.745 | 0.745 | 0.831 | 0.831 | 22 | 0.622 | 0.622 | 99 | 0.733 | 0.733 |
| **ropius0QA** | 186 | 0.731 | 0.731 | 0.826 | 0.826 | 22 | 0.537 | 0.537 | 99 | 0.718 | 0.718 |
| Robetta | 184 | 0.726 | 0.734 | 0.818 | 0.827 | 21 | 0.504 | 0.528 | 98 | 0.709 | 0.716 |
| IntFOLD6-TS | 184 | 0.679 | 0.687 | 0.785 | 0.793 | 21 | 0.387 | 0.405 | 98 | 0.647 | 0.654 |
| server999<sup>1</sup> | 173 | 0.678 | 0.729 | 0.783 | 0.841 | 20 | 0.432 | 0.476 | 92 | 0.650 | 0.699 |
| IntFOLD5-TS | 185 | 0.674 | 0.678 | 0.782 | 0.786 | 22 | 0.384 | 0.384 | 98 | 0.639 | 0.645 |
| IntFOLD3-TS | 186 | 0.660 | 0.660 | 0.770 | 0.770 | 22 | 0.328 | 0.328 | 99 | 0.629 | 0.629 |
| SWISS-MODEL | 185 | 0.649 | 0.652 | 0.755 | 0.759 | 21 | 0.252 | 0.264 | 99 | 0.613 | 0.613 |
| RaptorX | 159 | 0.583 | 0.682 | 0.665 | 0.778 | 19 | 0.358 | 0.414 | 81 | 0.531 | 0.649 |
| Phyre2 | 185 | 0.573 | 0.576 | 0.721 | 0.725 | 21 | 0.191 | 0.200 | 99 | 0.539 | 0.539 |
| IntFOLD4-TS | 158 | 0.571 | 0.672 | 0.659 | 0.776 | 17 | 0.300 | 0.388 | 89 | 0.576 | 0.641 |
| NaiveBLAST | 166 | 0.526 | 0.590 | 0.632 | 0.708 | 15 | 0.101 | 0.149 | 89 | 0.468 | 0.521 |
| PaFold | 114 | 0.443 | 0.723 | 0.497 | 0.812 | 13 | 0.364 | 0.616 | 66 | 0.475 | 0.713 |
| SPARKS-X | 93 | 0.292 | 0.583 | 0.351 | 0.703 | 14 | 0.179 | 0.281 | 40 | 0.220 | 0.545 |
| server99<sup>2</sup> | 38 | 0.147 | 0.720 | 0.158 | 0.773 | 1 | 0.019 | 0.417 | 27 | 0.193 | 0.709 |
| PRIMO | 44 | 0.142 | 0.602 | 0.163 | 0.690 | 4 | 0.050 | 0.277 | 31 | 0.181 | 0.579 |
| server102 | 41 | 0.136 | 0.618 | 0.167 | 0.759 | 3 | 0.033 | 0.242 | 23 | 0.130 | 0.558 |
| PRIMO_BST_CL | 40 | 0.134 | 0.623 | 0.152 | 0.709 | 3 | 0.043 | 0.315 | 26 | 0.148 | 0.565 |
| PRIMO_BST_3D | 37 | 0.114 | 0.575 | 0.134 | 0.673 | 3 | 0.032 | 0.236 | 27 | 0.150 | 0.549 |
| server100<sup>3</sup> | 26 | 0.110 | 0.785 | 0.115 | 0.822 | 1 | 0.019 | 0.417 | 18 | 0.143 | 0.786 |
| PRIMO_HHS_CL | 16 | 0.048 | 0.552 | 0.053 | 0.620 | 2 | 0.035 | 0.386 | 10 | 0.050 | 0.499 |
| PRIMO_HHS_3D | 15 | 0.043 | 0.530 | 0.046 | 0.567 | 3 | 0.036 | 0.265 | 9 | 0.051 | 0.559 |

## S2 Supplementary Methods

### S2.1 REDCNN training

Two datasets were prepared. Training was performed on one dataset, while the other served for validation of a trained instance of the REDCNN model. The diverse and independent datasets were obtained by applying these steps.

The initial set of sequences was first obtained from PDB sequences (02/2019) whose structures had been solved at a resolution of 2.5Å or less. These sequences were then clustered at 20% sequence identity using the blastclust program (Altschul *et al*., 1997) with soft masking of low complexity regions and the sequence-length coverage threshold set to 0.7. The representatives of the smallest clusters, classified in SCOPe database v2.07 (Chandonia *et al*., 2019), were split into two datasets so that the sequences of one dataset shared no common folds with the sequences of the other dataset. Random sampling from these two datasets separately yielded the training and validation datasets of predetermined sizes, 2001 and 200, respectively. The actual size of the training dataset was 1062 due to the minimum sequence length set to 128.

The training was performed on promages constructed for the sequences of the training dataset to predict distances between the CB (CA for Glycine) atoms observed in the corresponding protein structures at distances less than 64Å.

The REDCNN model was built using the Semantic segmentation suite (Seif, 2019), implemented in TensorFlow v1.15 (Abadi *et al*., 2016) and trained on an NVIDIA Tesla V100 GPU using a batch size of 4 randomly sampled 128 × 128 promage fragments and an RMSProp optimizer (Tieleman and Hinton, 2012) with a decay factor of 0.9 and a learning rate of 0.001, which was decreased to 0.0001 after the first 67% of training epochs. The objective function to optimize was a cross-entropy loss function.

Three independent training sessions were initiated. Three trained instances from the three sessions were selected based on manual inspection of predicted distances and the mean absolute error (MAE) between predicted distances and distances observed in all structures of the validation dataset (Figure S8). MAEs were separately calculated for all predicted distances (<64Å) and distances less than or equal to 8Å. Predictions for pairs of residues separated by less than four residues in the sequence were ignored.

We considered two approaches for predicting the distance between two residues. The distance with the highest probability corresponded to the predicted distance according to the first approach. And the predicted distance was the mean of the predicted probability distribution over distances according to the second approach. We used the first approach.

### S2.2 Effective number of sequences

The effective number of sequences (Marks *et al*., 2011) Neff in the MSA (Fig. 4, main text) is calculated using the neff program from the COMER2 software (Margelevičius, 2020). It is equal to

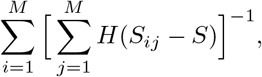

where *M* is the number of sequences in the MSA, *S_ij_* is the sequence identity between sequences *i* and *j* (normalized by the smaller sequence length), *S* = 62% is the sequence identity threshold, and *H*(·) is the unit step function.

**Figure S8.**
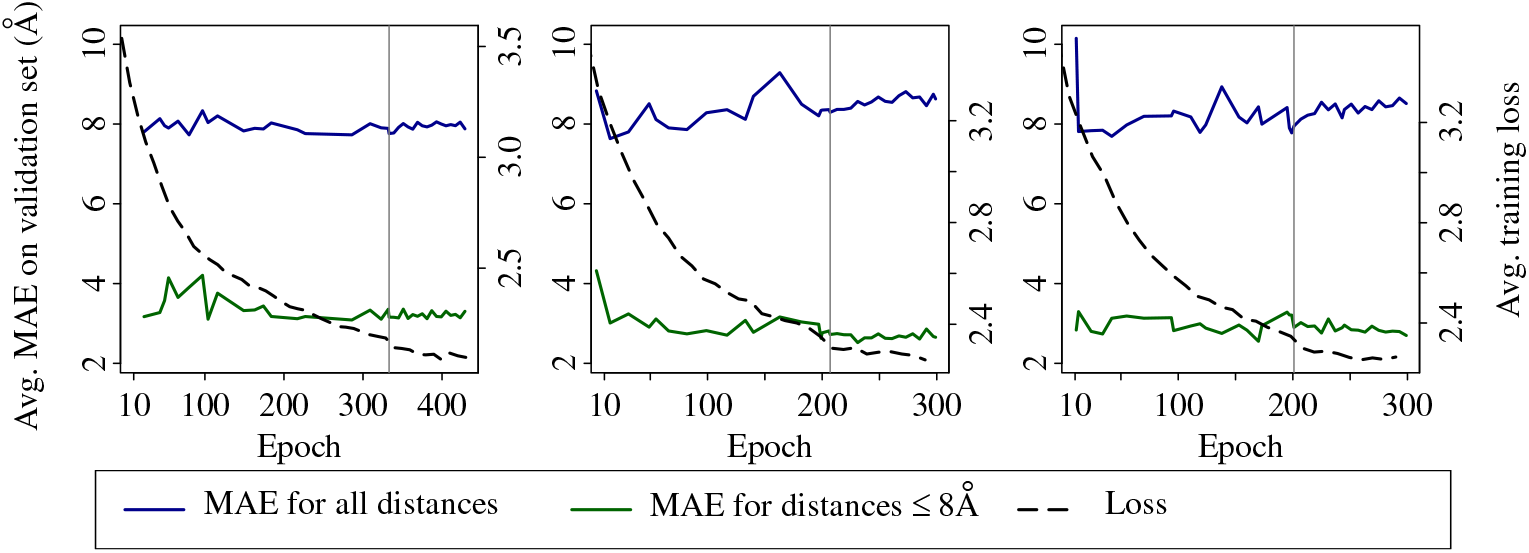
Average MAE on the validation dataset and average training loss versus training epoch for three independent runs of training. The standard deviation of MAE calculated for all predicted distances (<64Å; blue line) ranges from 6.4 to 8.7Å across all the runs for epochs greater than 20. It ranges from 4.0 to 5.5A for MAE calculated for distances predicted to be ≤8Å (green line). The straight vertical lines show the epochs at which three trained instances of the REDCNN model from different runs were selected for application.

